# Diel expression dynamics in filamentous cyanobacteria

**DOI:** 10.1101/2024.09.03.611090

**Authors:** Sarah J. Kennedy, Douglas D. Risser, Blair G. Paul

## Abstract

Filamentous cyanobacteria of the *Nostocaceae* family are able to differentiate into multicellular forms to adapt to environmental stresses, and members can establish symbiosis with various embryophytes. Representative laboratory strains are typically grown under continuous light to maintain stable metabolic conditions, however, this departure from a natural diel cycle can result in extended stress. Early genomic examination of *Nostoc punctiforme* suggests the genetic potential for a circadian clock, but we lack insight into global cellular dynamics through the natural diel cycle for this model organism. Here, we comprehensively assess changes in expression of core cellular processes and the mobilome of accessory genetic elements during diel growth of *N. punctiforme* PCC 73102. The primary transcriptome confirmed that multicellular cyanobacteria precisely coordinate photosynthesis and carbon assimilation for cell division during the day, while control of DNA recombination and repair appeared to be sequestered to darkness. Moreover, we expanded the known repertoire of light sensing proteins to uncover a putative regulator of circadian rhythm that itself exhibits striking oscillation between day-night expression. This was in sharp contrast to the arrhythmic pattern observed for a homolog of the canonical circadian regulator in unicellular cyanobacteria. Looking beyond cellular coordination of diel growth, we uncovered dynamic mobile elements, and notably, targeted hypermutation by retroelements that are likely maintained for conflict mitigation, which is crucial to a multicellular lifestyle.

## Introduction

Multicellular cyanobacteria, particularly those from taxonomic subsections IV and V, are thought to have evolved in several independent stages of life’s history, and occupy a vast range of environments (1)(2), where they adapt to variable stresses and serve as key drivers of ecosystem resilience. These microorganisms are able to differentiate several distinct cell types, including nitrogen-fixing heterocysts that shield the inside of the cell from oxygen (3), motile hormogonia that are thought to establish symbiosis with plant hosts (4), and desiccation-tolerant akinetes that develop and undergo dormancy in response to nutrient limitation or other stresses (5). *Nostoc punctiforme* strain PCC 73102 (ATCC 29133) is a model organism for studying development of these specialized cell types (6)(7)(8)(9).

Filamentous and multicellular cyanobacteria have been investigated, in large part, through experimental manipulation of model strains that represent members of the family *Nostocaceae* (4, 10). Growth experiments with *Nostoc punctiforme* PCC 73102, *Nostoc sp.* strain PCC 7120, and other model representatives are typically carried out under continuous light (i.e. 24-hours constant illumination), as opposed to mimicking the natural light-dark diel cycle. Constant light incubations have been used in the past for examining the ability of various cyanobacteria to fix nitrogen (11), maintain photosystem regulation (12), establish circadian rhythm (13, 14), and undergo chromosome supercoiling (15). Yet, continuous light exposure has been linked to oxidative stress and upregulation of protective response pathways (12). The departure from the natural diel cycle to continuous light induces global transcriptional changes in cyanobacteria, complicating the interpretation of gene expression due to stress-related perturbations and adaptive responses. Moreover, transcriptomic studies with *N. punctiforme* have not yet addressed the natural cellular response to a diel cycle and in turn, whether the organisms can establish circadian rhythm from long-maintained cultures. As such, we lack a complete view of global regulation during natural growth of model filamentous cyanobacteria.

The genomes of multicellular cyanobacteria are notably expansive, especially when compared with some unicellular counterparts that maintain smaller genomes (16, 17), which has been attributed to rampant duplication and horizontal gene exchange. Of particular note are the functional roles and mechanisms underlying expansion of diverse mobile elements and variable repertoires of protein kinases in multicellular cyanobacteria, which remain to be characterized. Moreover, the balance between maintenance and differential expression for multiple gene copies raises questions on the evolutionary fate of these expanding loci (18).

In this study, we established diurnal entrainment of *Nostoc punctiforme* and generated a global gene expression dataset, from which we systematically uncovered dynamic cellular processes that oscillate through the light-dark cycle. This study provides a new framework and primary resource for interpreting dynamic gene expression in filamentous cyanobacteria undergoing diel responses to light, while establishing circadian rhythm.

## Results and Discussion

To uncover dynamic cell processes during diel growth in *Nostoc punctiforme*, we carried out a large-scale growth and multi-omics experiment, comprising genome re-sequencing and 36 transcriptomic libraries. Cultures were grown on a light-dark (LD) diurnal cycle (i.e. 12 hours light followed by 12 hours dark), and four distinct stages were targeted to generate transcriptomes (**Figure 1A**). In the first stage, two timepoints were harvested after 36 and 48 hours – referred to here as “Pre-Dawn0” and “Pre-Dusk0”, respectively. Next, the second and third stages were intended to observe early versus late diurnal entrainment, separated by three days. To this end, both the early and late entrainment stages were sampled at 6-hour intervals encompassing: i) mid-phase dark, ii) 30-minutes pre-dawn, iii) mid-phase light, iv) 30-minutes pre-dusk. In the fourth stage, diel cultures grown for 8 days were incubated for a final 24 hours in either i) continuous light (LL) or ii) continuous dark (DD). Global RNA expression was assessed for all four sampling stages, comprising 12 time points measured in triplicate (**Figure 1B**).

**Figure 1.**
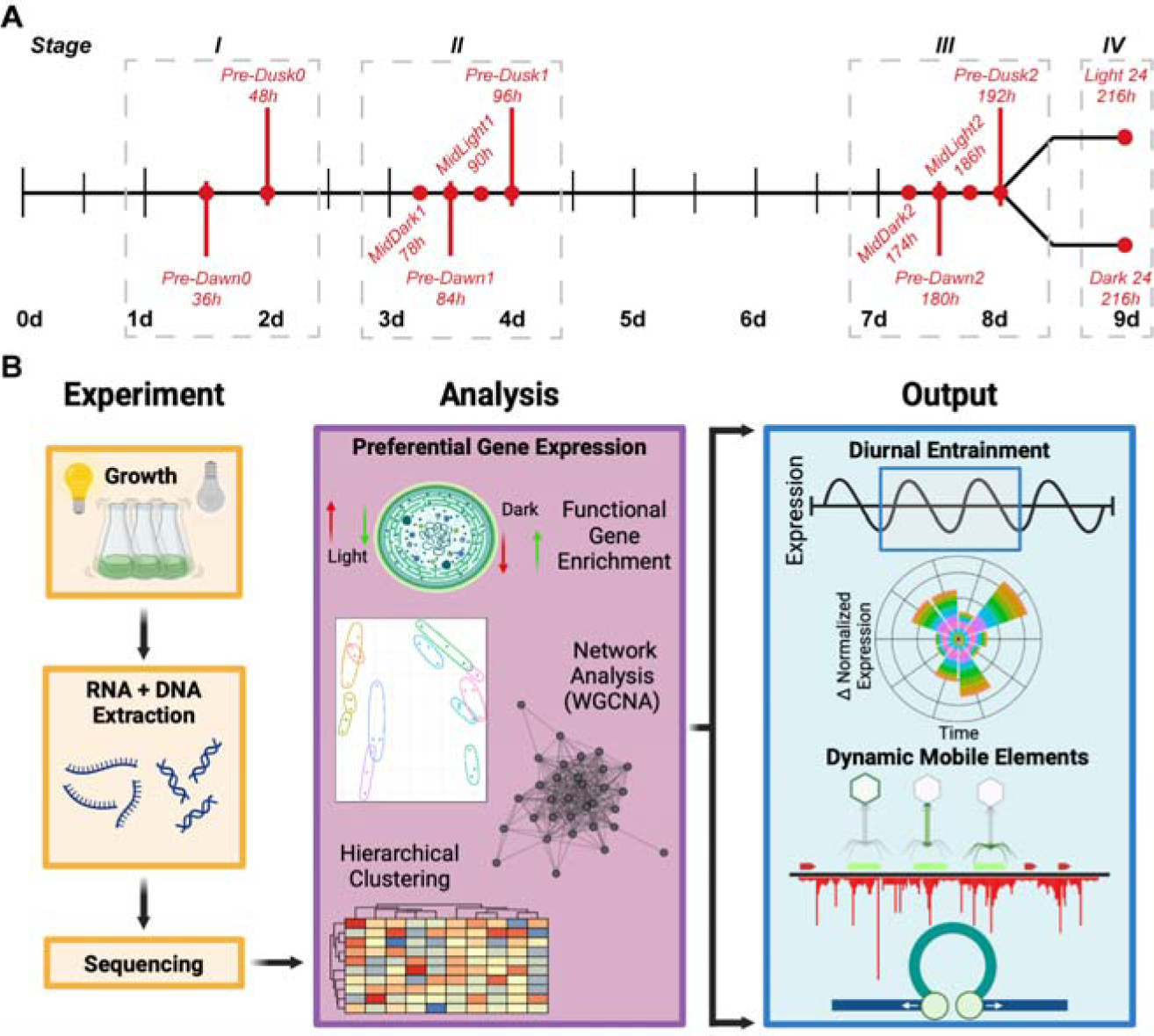
Overview of experimental design and analytical workflow. (**A**) Schematic of the experimental design and timeline. Sampling spanned 9 days, including 12 timepoints to capture pre-dawn, pre-dusk, mid-dark, and mid-light phases. Four sampling stages are indicated (I-IV) that encompassed early diurnal entrainment, late diurnal entrainment, and reversion to continuous light or dark. Individual timepoints are highlighted with the respective light or dark state. **(B)** Workflow illustrating the experimental design and analysis. Experimental steps include diel growth, RNA and DNA extraction, and sequencing. The preferential gene expression workflow included functional gene enrichment, statistical analyses, weighted gene co-expression network analysis (WGCNA), and hierarchical clustering. The workflow outputs revealed gene expression dynamics during diurnal entrainment, and active mobile elements, such as phage and transposons. Images created with BioRender.

From a global perspective, the individual transcriptomes (n = 36) are most similar among replicates (**Supplemental Figure S1**), then primarily separate according to growth in the light vs dark (**Supplemental Figure S2**). Broadly, transcriptomes from the light phases (Pre-Dusk, MidLight, Light24) contrast with those from dark phases (Pre-Dawn, MidDark, Dark24), while within-phase clusters of individual samples differentiate among the respective timepoints. Moreover, multiple distinct temporal clusters emerge from this global analysis: i) initial dark, ii) initial light, iii) mid-light, iv) pre-dusk, v) mid-dark, vi) pre-dawn, vii) 24-hr-post in dark, viii) 24-hr-post in light.

### Global response to light availability

To assess diurnal gene expression in *N. punctiforme*, we first looked at the core periods of our time series, which encompassed two full 24-hr LD cycles (i.e. Stages II and III; **Figure 1A**). Hierarchical clustering of transcriptomes for these core timepoints revealed two overarching partitions (**Figure 2A**), apparently corresponding to the light/dark state. By assessing expression variation across groups of “Light” (Mid-Light and Pre-Dusk) and “Dark” (Mid-Dark and Pre-Dawn) timepoints, we found 200 genes and predicted RNAs that had significantly different expression profiles across the core diurnal entrainment. Among this subset, 117 genes/RNAs were preferentially expressed within mid-light and pre-dusk timepoints (**Figure 2B**). This set of genes was associated with diverse metabolic pathways; broadly, these include general energy, carbohydrate, amino acid, and other core pathways – including metabolisms of lipids, nucleotides, and cofactors/vitamins. Notably, preferential gene expression was most similar between adjacent timepoints (e.g., MidLight1 and Pre-Dusk1), rather than by condition (e.g., MidLight1 and MidLight2; Pre-Dusk1 and Pre-Dusk2). In contrast, 83 variable genes and predicted RNAs were preferentially expressed within dark-associated conditions, with the majority of gene functions remaining poorly characterized (**Figure 2C**).

**Figure 2.**
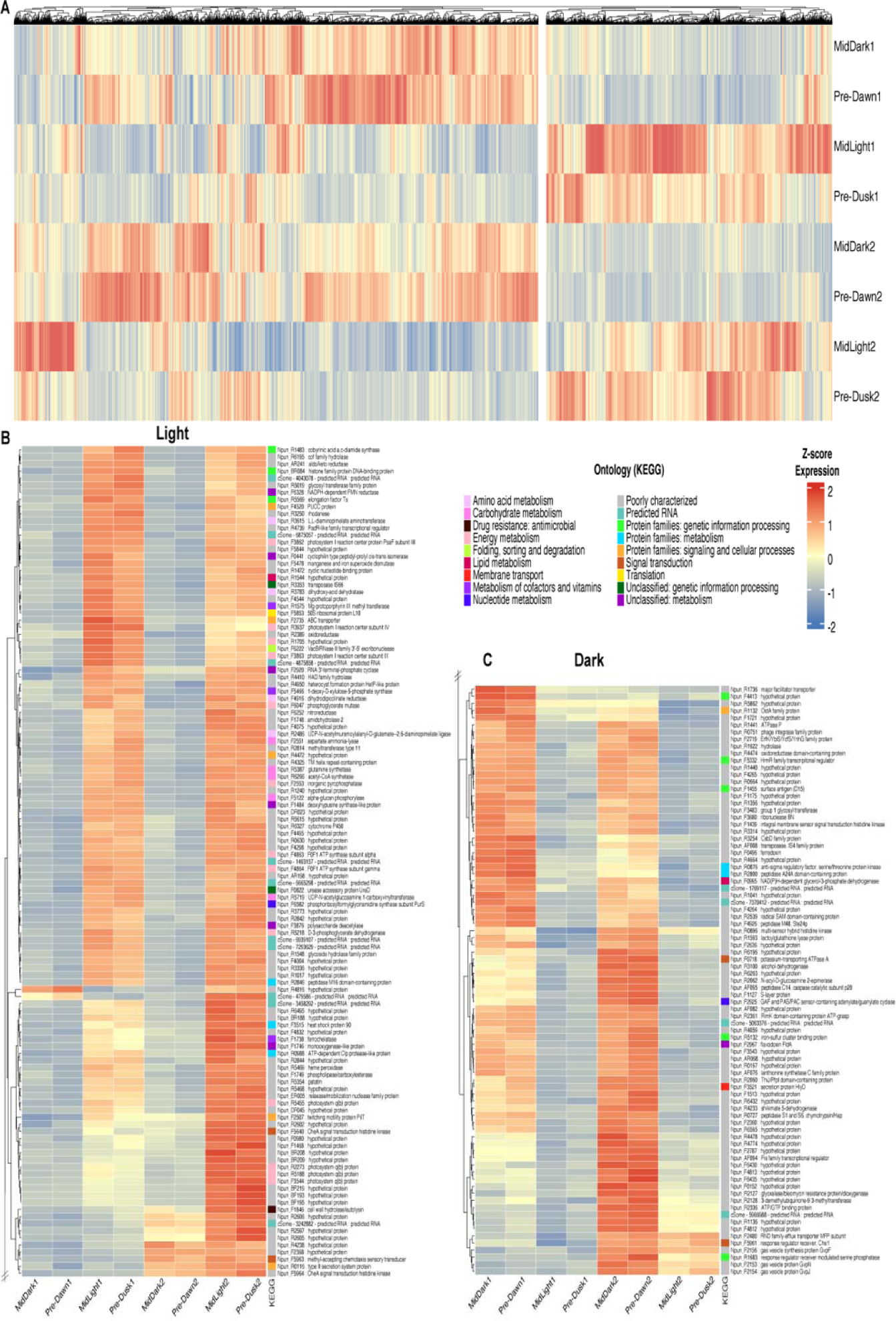
Preferential gene expression across diurnal timepoints. (**A**) Heatmap of gene expression from sampling stages II and III (MidDark1, Pre-Dawn1, MidLight1, Pre-Dusk1, MidDark2, Pre-Dawn2, MidLight2, and Pre-Dusk2). The data were hierarchically clustered using Euclidean distance and complete linkage. Each dendrogram branch represents a gene, with z-score normalized expression levels (i.e., blue indicates downregulation and red indicates upregulation). **(B)** Light-significant and **(C)** dark-significant preferential gene expression (BH-adjusted, ANOVA) during light and dark timepoints. KEGG annotations are represented with distinct colors corresponding to their functional categories.

Looking beyond the core LD timepoints, we next clustered the full experimental transcriptional dataset, which revealed “light-’’ and “dark-’’ associated expressions (**Supplemental Figure S3**). Specifically, we carried out weighted gene co-expression network analysis (WGCNA) for all 12 timepoints, resulting in gene modules that display co-regulation and thus may underlie common cellular functions (**Supplemental Figure S4A**). Importantly, the WGCNA modules were analyzed and grouped based on their correlations with timepoints (**Supplemental Figure S4B**), which were used to determine preferential expression.

Together, all modules that were associated with light comprised 1,035 genes, whereas the dark-associated modules comprised 776 genes. Strikingly, most of the light-associated genes were assigned to Light24 modules (507 genes), followed by Pre-Dusk (326 genes) and MidLight (202 genes) modules. Conversely, the majority of dark-associated co-expression was identified within Pre-Dawn (483 genes), followed by Dark24 (164 genes) and MidDark (129 genes) modules.

### Light-associated co-expression of metabolic pathways and motility systems

Having conducted separate analyses of global expression clustering (hierarchical), variation (ANOVA), and correlation (WGCNA), we next used the combined output of associated genes as input for functional enrichment analysis (**Supplemental Figure S3**). Functional gene set enrichment analysis (GSEA) revealed light-associated convergence of biosynthetic pathways, including synthesis of peptidoglycans, O-antigen nucleotide sugars, amino sugars, and other nucleotide sugars (**Figure 3**). Unsurprisingly, photosynthesis was the most significantly enriched pathway within light-available transcriptomes (particularly within Pre-Dusk and Mid-Light, **Figure 3A**). Interestingly, peptidoglycan biosynthesis is also enriched within Pre-Dusk and Mid-Light transcriptomes. While several peptidoglycan biosynthetic enzymes (and thus their intermediate/precursor metabolites) are shared amongst other light-enriched sugar metabolic pathways (**Supplemental Figure S5**), this notable convergence of co-expression (**Figure 3B** and **Supplemental Figure S6**) suggests concerted regulation within diurnally-entrained *N. punctiforme*.

**Figure 3.**
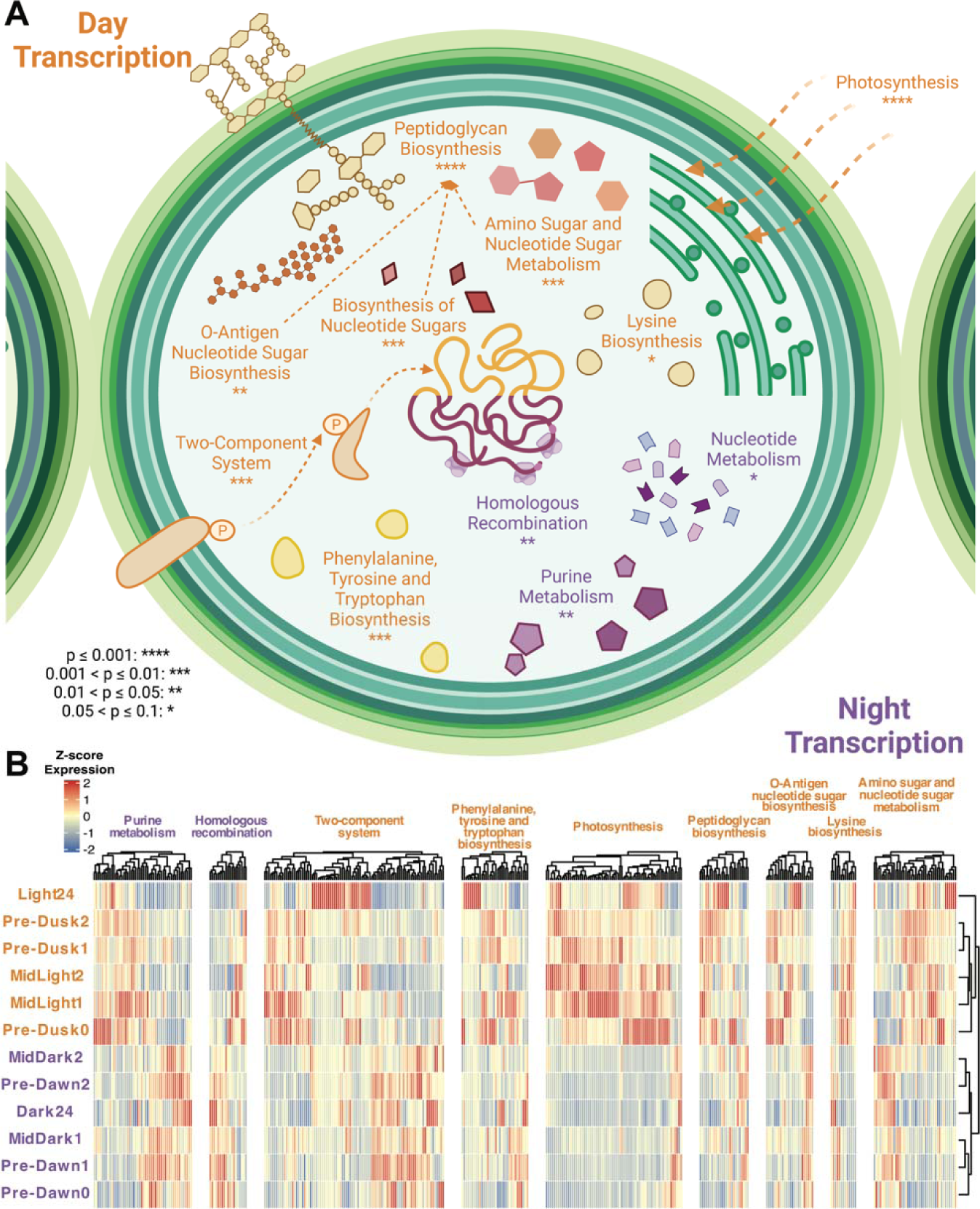
Gene expression and pathway enrichment analyses of diel transcription. (**A**) Cell process diagram depicting enriched pathways associated with day (orange) and night (purple) transcription. Significance levels (hypergeometric test) are indicated as follows: ****p ≤ 0.001, ***0.001 < p ≤ 0.01, **0.01 < p ≤ 0.05, *0.05 < p ≤ 0.1. **(B)** Heatmap of hierarchically-clustered gene expression profiles (Euclidean distance, complete linkage) within enriched pathways from the time-course experiment. Z-score normalized expression levels are color-coded, with red indicating upregulation, and blue indicating downregulation.

Insights into diel *Synechococcus elongatus* growth studies reveal the importance of glycogen cycling (19) for continued light-dark-cycle survival (20), however, closer examination of our transcriptional time-course data reveals the specific enzymatic paths linking metabolism with cell wall biosynthesis. Notably, our dataset reveals enrichment of enzymatic pathways intertwining glycolysis/gluconeogenesis intermediates (specifically, glucose-6-phosphate (Glc-6P) and fructose-6-phosphate (Fru-6P)) with other sugar metabolic pathways, culminating in the necessary precursors for peptidoglycan biosynthesis (**Supplemental Figure S5**).

While these specific enzymes are shared amongst pathways that apparently respond to a diel cycle (O-antigen nucleotide sugars; amino sugars; and nucleotide sugars), two genes (*nagB,* Npun_R1302; *glmU,* Npun_F0907) demonstrate correlated expression with multiple key peptidoglycan biosynthetic genes (**Supplemental Table S1**). Although highly correlated with only two peptidoglycan biosynthetic genes, *nagB* (catalyzing conversion of glucosamine 6-phosphate (GlcN-6P) to Fru-6P) expression demonstrates potential co-expression with both a low molecular weight PBP (LMW PBP, Npun_R1733, ∼0.87) and an isoprenyl transferase (*uppS,* Npun_R1952, ∼0.76). Conversely, *glmU* (responsible for catalysis of the last two steps of *de novo* biosynthesis of peptidoglycan precursor UDP-N-acetylglucosamine (UDP-GlcNAc)) expression correlates with five peptidoglycan biosynthetic enzymes. These include the genes responsible for the first committed step of peptidoglycan biosynthesis (*murA*, Npun_R5719, ∼0.86), precursor transport/insertion into the cell wall (*bacA*, Npun_R4507, ∼0.83), and crosslinking (LMW PBP, Npun_R2557, ∼0.83; Class B PBP, Npun_F4453, ∼0.79; Class B PBP, Npun_F0907, ∼0.77).

Critical cell division genes (*ftsE, minC, ftsK, cdv1,* and *minD*) are highly correlated (>0.90) with several direct peptidoglycan synthetic genes (class B PBP, *murE,* LMW PBP) and indirectly synthetic/convergent metabolic genes (*ddl, glmU;* **Supplemental Table S2**). Taken together, the cellular division-associated genes appear correlated by expression in the LD cycle. Tight co-expression of sugar metabolism, peptidoglycan biosynthesis, and cellular division suggests that cell wall growth and development follows precise timing mechanisms linked to the LD cycle, as previously demonstrated by circadian clock-gated cell division in *S. elongatus* (*21*).

These transcriptional observations are supported by metabolomic studies in diurnally-entrained *S. elongatus,* wherein daytime accumulation of various sugar metabolites is coordinated with increased rates of cellular growth (22), implicating carbon assimilation to the cell wall. Additionally, diel random bar-code transposon-site sequencing (RB-TNseq) with *S. elongatus* determined that these processes were vital during the LD growth versus continuous light (23), supporting our observations of carbon metabolism and cell wall synthesis converging with the LD cycle.

We next examined other functions not clearly tied to metabolism, which have preferential expression in the light. Notably, a subset of two-component system genes was found with high Light24 expression (**Supplemental Figure S7**). Closer inspection reveals that the majority of these preferentially expressed genes belong to chemotactic systems specifically expressed within developing hormogonia (24, 25). This preferential expression implies light or LD cycle influence on hormogonium development. Indeed, when genes with experimentally validated functions in hormogonia (**Supplemental Table S6**) are examined for preferential expression across the time-course series, the majority reveal significant upregulation within Light24 (**Supplemental Figure S8**). Pronounced Light24 expression could indicate that motile hormogonium development and underlying transcriptional cascades are activated in response to constant light.

### Nucleotide metabolism and DNA repair during darkness

Although most genes that were preferentially expressed in periods of dark remain poorly characterized, we were able to identify several functions that were significantly enriched and co-expressed in these timepoints. Elevated expression in the dark (mid-dark and/or pre-dawn) was most associated with purine metabolism, homologous recombination, and nucleotide metabolism (**Figure 3A**). Notably, these functions were most enriched in both Pre-Dawn1 and Pre-Dawn2 timepoints (**Figure 3B** and **Supplemental Figure S7**). By contrast, genes linked to these, or other core cellular processes were not more highly expressed in any mid-dark timepoints. This finding raised the question of why pre-dawn timing of expression might be critical for certain processes.

Prior studies have shown that circadian rhythm regulates the storage and conversion of sugars in *S. elongatus* (*26*), where specifically, RpaA regulates carbohydrate storage during the day and energy conversion at night. Separately, the transcriptional regulator cyAbrB was found to control the day-night shift in carbohydrate metabolism in *Synechocystis sp.* PCC 6803 (27). Transcriptomes of *N. punctiforme* imply that during darkness, stored sugars are primarily used to generate purines and nucleotide pools. The genes enriched during darkness are central to the pathways that convert nucleoside monophospates to nucleotide bases (**Supplemental Figure S9**).

It remains unclear whether homologous recombination genes are directly controlled by circadian regulators or respond independently to darkness. Transformation has been shown to coincide with the onset of darkness in *Synechococcus elongatus* (*28*). Taton et al. demonstrated that the circadian clock regulates natural competence, with type IV pilus machinery (T4PM) biogenesis genes expressed in the morning and T4P transformation genes maximally expressed during dusk (29). Taken together, these findings suggest a possible connection between circadian regulation and DNA repair mechanisms.

We next looked more closely at the individual genes that comprise the core cellular functions associated with pre-dawn. In particular, pronounced expression of *recR, recG, and recD* (Npun_F4120, Npun_R5567, Npun_AR019) was observed during pre-dawn timepoints (**Supplemental Figure S7**). RecD is the only component of the bacterial RecBCD complex encoded in the genome of *N. punctiforme*, and notably, it is a plasmid-encoded helicase. RecG is a multifunctional ATP-dependent DNA helicase, which is thought to contribute to chromosome repair, acting synergistically with the RuvABC complex (30). Interestingly, the diurnal transcriptome of *N. punctiforme* revealed this singular protein was enriched during the pre-dawn timepoint, which raises questions on the specific timing of DNA repair in filamentous cyanobacteria. The repair proteins were previously shown to be duplicated and highly conserved in several cyanobacterial genomes (31).

Two of six total proteins that can be components of the DpoIII complex were enriched in the pre-dawn transcriptome (**Supplemental Figure S7**): gamma/tau complex (Npun_F4123), and the alpha subunit (Npun_F5684). The gamma/tau subunit is predicted to promote processivity, through loading the beta clamp onto DNA (32). And moreover, the alpha subunit holds the catalytic core of DpolII, wherein DNA synthesis proceeds (33). Taken together with the diel expression of repair proteins, these findings suggest that filamentous cyanobacteria may sequester chromosomal maintenance during darkness. Furthermore, given the necessity for damaged DNA to be repaired for efficient replication by DpoIII, we speculate that RecG-based repair may be timed synergistically with DpoIII expression. Indeed, the gamma/tau complex expression correlated with *recR* (*r =* 0.63), *recG* (*r* = 0.91), and *recD* (*r* = 0.90) expressions. By contrast, DpoIII alpha subunit expression negatively correlates with that of *recR* (*r =* –0.78), *recG* (*r* = –0.81), and *recD* (*r =* –0.75).

We next examined gene expression of repair proteins during the Dark-24 timepoint, which represents a perturbation of the diel cycle. Notably, a cluster of four DNA processing and repair genes were disproportionately expressed following this perturbation, comprising Dpo III α and δ, ssDNA exonuclease RecJ, and repair protein RecO (**Supplemental Figure S7**). Importantly, expression of these genes during Dark-24 was in sharp contrast with their comparatively low expression during the Light-24 perturbation, suggesting a specific response to darkness. Finally, many genes that were clustered with highest expression under dark conditions remain hypothetical/uncharacterized. Thus, it is unclear whether other cellular pathways are programmed to be upregulated in the dark.

### Transcription of circadian and clock-controlled proteins

While the global transcriptome appeared to partition in apparent response to light, circadian clock-controlled expression is most prominent during the pre-dawn/pre-dusk transitional timepoints. Cyanobacterial circadian rhythms are driven by proteinaceous clock-controlled proteins and post-transcriptional regulation (34), as demonstrated within the non-cyclical expression of *kaiABC* core circadian genes (**Supplemental Figure S10A**). Notably, however, circadian clock-associated regulators (*sasA, cikA, rpaA,* and *rpaB;* **Supplemental Figure S10B**) exhibit cyclical expression peaking within pre-dawn transcriptomes. Importantly, the GAF sensor CikA plays an intermediary role linking the LD cycle with the circadian clock. Notably, while the KEGG-identified CikA (Npun_F1000) demonstrates high amino acid similarity (50.50%; **Supplemental Table S3**) to the canonical *Synechococcus elongatus* CikA, no pronounced expression is observed across the time course series (**Supplemental Figure S10B**).

In order to explore alternative kinases that may fulfill the CikA role within *N. punctiforme*, we identified twelve *cikA*-like genes encoded throughout the genome based on sequence similarity to cyanobacterial proteins containing CikA-like GAF and histidine kinase domains (**Supplemental Figure S11A; Supplemental Table S4**). While these genes contain sequence similarity to canonical CikA proteins, we sought to assess expression across the global transcriptome for functional alternative kinases that may fulfill the CikA role in *N. punctiforme.* Revisiting the Pearson correlation analysis, we uncovered a subset of genes associated (r ≥ 0.75) with cell division, peptidoglycan biosynthesis, and intermediary genes linking central carbon metabolism with peptidoglycan biosynthesis (**Supplemental Table S2**).

Among these, a single GAF-like histidine kinase (Npun_R5149) correlated with five genes of interest: Npun_F0907 (*glmU;* precursor to peptidoglycan biosynthesis); Npun_F2411 (*murG*; peptidoglycan biosynthesis); Npun_F4453 (Class B PBP; cell wall synthesis); Npun_R4507 (*bacA*; cell wall lipid modification); and Npun_R4933 (*cdv1*; cell division). Notably, this putative CikA-like gene exhibits cyclical expression most pronounced during pre-dusk and pre-dawn timepoints (**Supplemental Figure S11B**).

Interestingly, another *N. punctiforme cikA*-like gene (Npun_R2903) shows apparently cyclical expression. Despite not being as highly expressed as Npun_R5149, Npun_R2903 had consistently higher expression than the canonical *cikA* across timepoints (Npun_F1000; **Supplemental Figure S11B**). Moreover, hierarchical clustering of the correlations among associated genes described above (**Supplemental Figure S11C**) revealed similarity to Npun_R5149 and canonical *cikA* (Npun_F1000). Both alternative CikA-like, GAF-domain-containing histidine kinases demonstrate low amino acid similarity to canonical *S. elongatus* CikA (Npun_R5149 29.60%; Npun_R2903 26.10%; **Supplemental Table S3**) and cyanobacterial CikA-like proteins (Npun_R5149 26.90%; Npun_R2904 29.90%; **Supplemental Table S4**). Despite low sequence similarities to the annotated CikA locus (Npun_F1000), these other GAF-like histidine kinases (Npun_R5149 and Npun_R2903) demonstrate higher and more cyclical/dynamic expression as compared to the annotated *cikA* locus, perhaps fulfilling the role of a regulatory link between the LD cycle and the circadian clock. Indeed, when exploring the transcriptome for potentially co-expressed genes, Npun_R5149 correlates with the greatest number of genes as compared to Npun_R2903 and Npun_F1000 (**Supplemental Figure S11D**).

Critically, canonical *cikA* (Npun_F1000) only significantly (≥0.75) correlates with one gene implicated in the LD cell growth/division geneset (Npun_F5950, *ylmG:* nucleoid distribution), which contrasts the other more intercorrelated CikA-like histidine kinases (**Supplemental Table S5**).

We next sought to further examine the domain architecture and phylogeny for this GAF-like kinase protein (**Supplemental Figure S11E**). It appears that Npun_R5149 and homologs from other filamentous cyanobacteria may have evolved from CikA of unicellular cyanobacteria, to function in establishing the circadian clock for filamentous cyanobacteria. Most notably, whereas all CikA-like homologs maintain one histidine kinase and ATPase domain, in stark contrast to Npun_F1000, other loci encode an array of multiple GAF domains, duplicated PAS and PAC signal sensors, and one or more receiver domains. The GAF-containing histidine kinases (GAF-HK) of *N. punctiforme* show architectural complexity through fusion of extracellular sensory domains (CHASE-type), histidine kinase phosphotransfer domains (HPT), and incorporation of transmembrane sites. The subset of highly expressed and dynamic GAF-HK (Npun_R5149, Npun_F6362, Npun_R6149, Npun_R2903) has a complex array of sensor and receiver domains, when compared with Npun_F1000 and the CikA of *S. elongatus* (Synpcc7942_0644).

Although circadian mechanisms have been explored extensively within *S. elongatus* systems (35)-(36), and physiological evidence of circadian rhythm is lacking for *N. punctiforme* (*37*), recent gene reporter experiments demonstrated circadian clock activity in the closely-related filamentous cyanobacterium, *Anabaena* sp. PCC 7120 (elsewhere known as *Nostoc* sp. PCC 7120; (36)). Circadian clock-controlled proteins identified within this similarly multicellular and filamentous model organism share homology to several *N. punctiforme* genes with apparent clock-influenced expression (**Supplemental Figure S12**). While some of these homologs show transcriptional oscillation, separate studies suggest that circadian transcripts can be uncoupled from circadian protein activity, and vice versa (38) (35) (34).

### Mobile genetic elements are dynamically expressed

*Nostoc punctiforme* has been developed for several decades as a model organism for studying symbiosis and cellular differentiation in filamentous cyanobacteria. Yet relatively little is known about the role of mobile genetic elements (MGE) in this organism. Many MGE are well-characterized in pathobionts of humans and animals, where these systems play an important role in virulence (39–41). Moreover, in environmental and non-pathogenic microorganisms, the same MGE can often be identified via bioinformatic surveys, however their functional significance and genetic activity are overlooked. We thus took advantage of our robust temporal dataset to explore the dynamic subset of MGE in this model of filamentous cyanobacteria. Hierarchical clustering analysis of the global dataset revealed several groups of genes with similar expression profiles across timepoints, including IS4 transposase genes, and two putative prophage elements (**Supplemental Figure S13**). Interestingly, a group of co-localized phage-like genes showed pronounced expression in the MidDark2 timepoint (Npun_F1091 – Npun_F1118), including five putative tail fiber proteins (**Figure 4A**).

**Figure 4.**
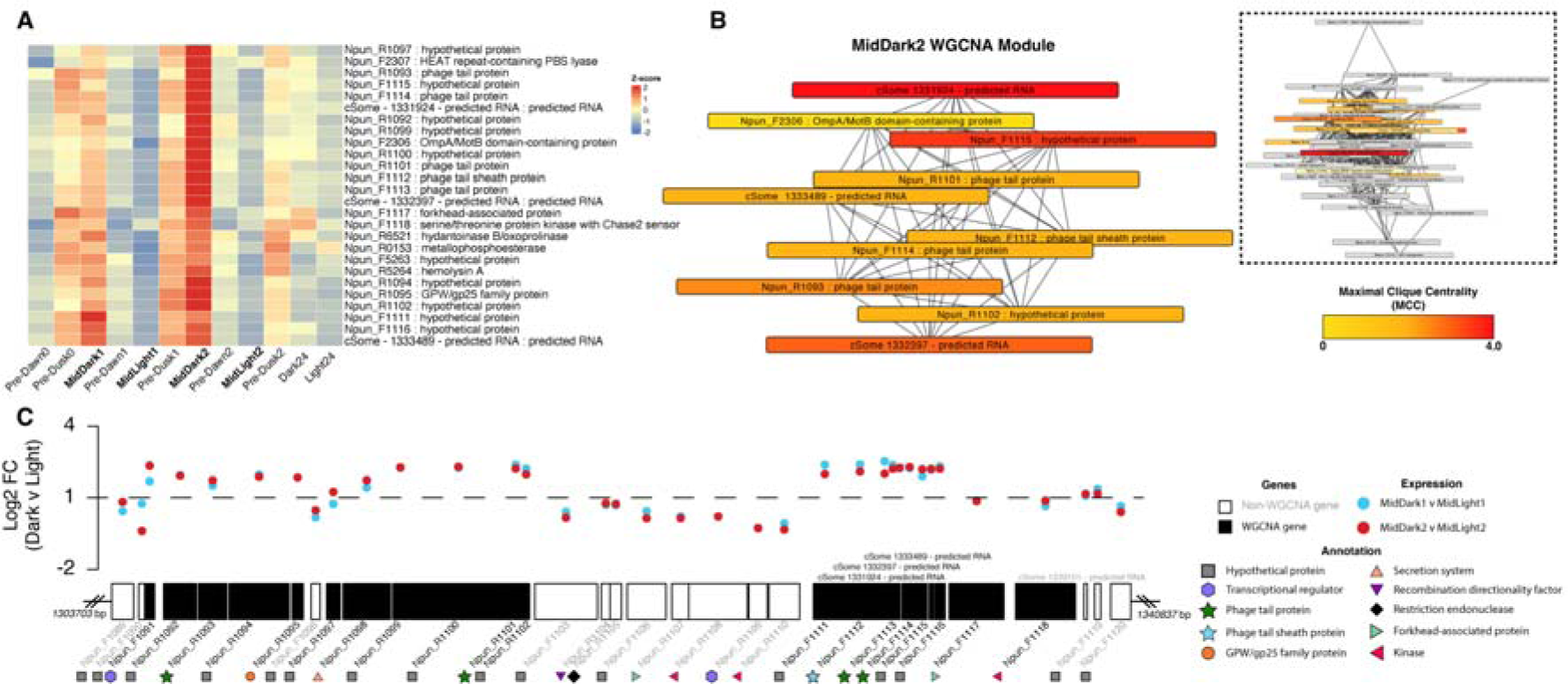
Analysis of MidDark2-associated putative prophage. (**A**) Heatmap of expression data from hierarchical clustering of the entire transcriptome, highlighting the MidDark2 phage gene cluster. Each gene’s expression is shown across each timepoint, with Z-score normalized expression (downregulated = blue, upregulated = red). **(B)** The most associated weighted gene co-expression network analysis (WGCNA) module for MidDark2, including phage genes, distilled into the top 10 Maximal Clique Centrality (MCC) nodes. The MCC scores indicating the centrality and importance of genes within the network are represented by a color gradient from low (yellow) to high (red). **(C)** Chromosomal region ranging from 1,303,703 bp to 1,340,837 bp encoding for phage-associated genes (black shading) and non-phage associated genes (white). Above the loci, log2 fold change (Log2 FC) is plotted for genes comparing MidDark1 vs. MidLight1 (blue) and MidDark2 vs. MidLight2 (red) timepoints. Annotations, where available, are provided for each locus.

We next used network analysis to uncover additional host genes that correlate with the putative prophage element. Notably, the prophage locus associated with the MidDark2 timepoint was central to a weighted co-expressed gene network (**Figure 4B**). This network was exclusively correlated with MidDark2, and stands out among other networks within the same timepoint (**Supplemental Figure S4**). Crucially, WGCNA confirmed that all of the phage tail genes are indeed co-expressed and revealed their broader network of associated genes; within this module, the ten genes that represent the maximum clique centrality include four of the five phage structural genes (**Figure 4B**). While this network encompasses a set of co-localized phage genes, other proximal genes show distinct expression profiles (**Figure 4C**).

Within this gene neighborhood, we uncovered two apparent modes of expression, distinguished by putative structural genes, versus genes with predicted recombination and regulation functions. The putative phage structural genes showed elevated expression (Npun_F1091 – Npun_R1102; Npun_F1111 – Npun_F1116) in MidDark2, whereas non-structural genes (Npun_F1103 – Npun_R1110) were notably absent in the MidDark2 co-expression network. We hypothesize that the network of genes expressed during MidDark2 – including phage tail, sheath, and a putative head morphogenesis GpW protein – are associated with virion assembly. These findings suggest that the temporal transcriptome captured a switching event, wherein a temperate virus may have been induced into its productive lytic cycle.

To assess whether this observation represents a stochastic event, or a persistent phenomenon, we carried out an additional growth experiment designed to target the same diel intervals (**Supplemental Figure S14**). Quantitative amplification of the locus, Npun_F1112 (phage tail sheath protein), confirmed that the putative phage gene was most expressed in the MidDark timepoint in both our original growth experiment and in the repeated experiment. Although median expression was highest in MidDark during the follow-up growth experiment, we did observe wide variation across replicates of the Pre-Dawn timepoint, wherein one replicate appeared to have higher expression than that of MidDark. Nonetheless, this independent assay supports the hypothesis put forth that structural genes found in a putative prophage element are preferentially expressed in the dark. Still, despite expression of phage structural genes suggesting that the elements produce infective viral particles, they may instead represent replicative and integrative transposons that resemble MGE from other cyanobacterial mobilomes (42).

While cyanophage of globally abundant marine unicellular phytoplankton have been described in great detail, the viruses of filamentous cyanobacteria remain relatively understudied. A collection of phage isolates from environmental blooms of *Nostoc* were previously described (43, 44), and more recently, three phage genomes were sequenced and contextualized (45). The putative temperate phage uncovered in this study is distinct and may represent a useful model to interrogate the host-phage dynamics and the mechanisms underpinning lysogeny in filamentous cyanobacteria.

### Hypermutation of conflict mitigation systems

Diversity-generating retroelements (DGRs) are domesticated retrotransposons that drive targeted protein evolution and were recently highlighted as prevalent within genomes of filamentous cyanobacteria. In most DGRs, a genomic locus encodes essential features, comprising a reverse transcriptase (RT) that complexes with an accessory variability determinant protein (Avd), template RNA repeat (TR) and cognate variable repeat (VR) coding site(s), which differ from the TR at adenines. The putative DGR of *N. punctiforme* was previously described and has two target genes (17), yet adenine mutagenesis has not been directly observed in these or other cyanobacterial cultures. The essential DGR components are all expressed (**Figure 5A**), but one target gene appears disproportionately expressed in comparison with the second target. The known diversifying machinery, comprising RT and putative Avd, are consistently expressed across timepoints, though to a lesser degree than the neighboring genes.

**Figure 5.**
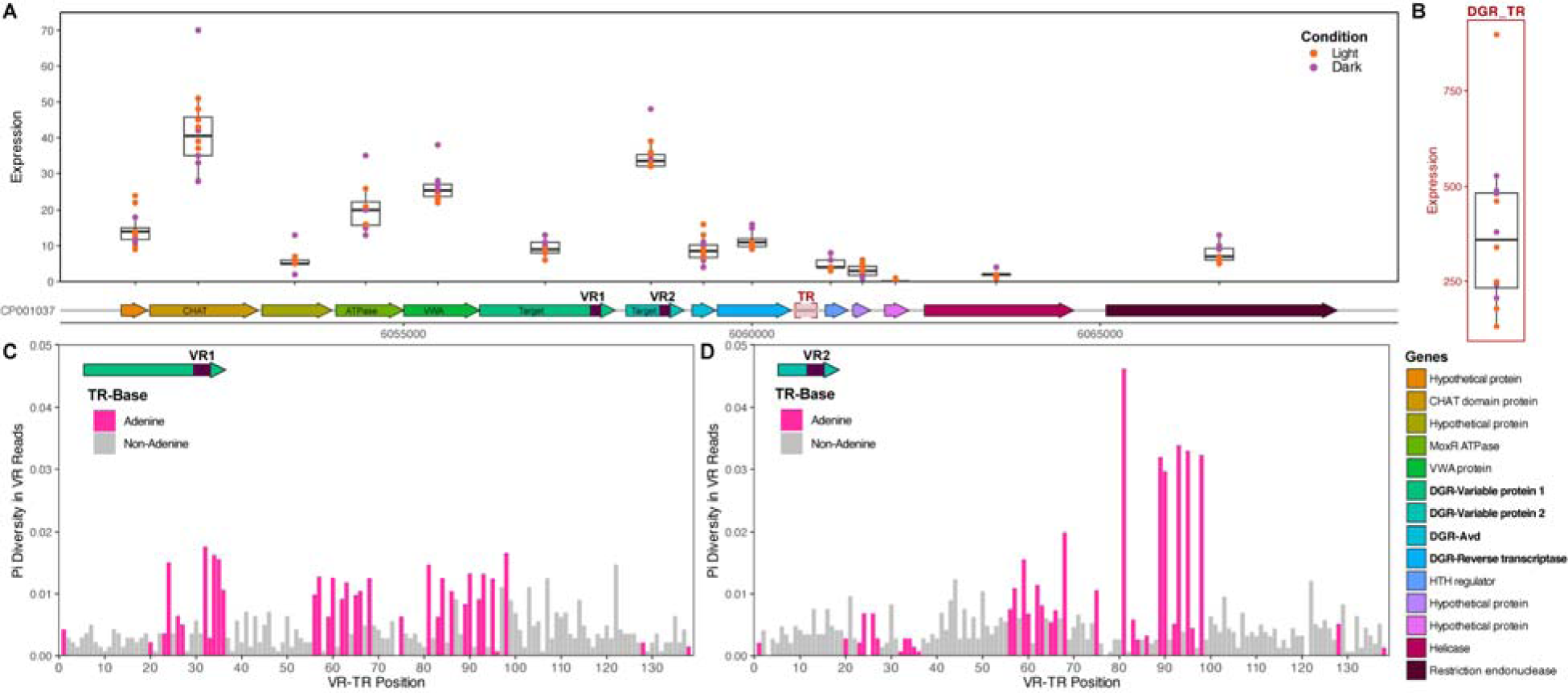
Diversity-generating retroelement (DGR) analysis. (**A**) Chromosomal region surrounding DGR locus, annotated with color-coded gene functions. Box plots represent the expression levels across 12 different timepoints, with each box plot displaying the median, quartiles, and range of expression values under light (orange) and dark (purple) conditions. Purple shaded boxes indicate the variable regions (VR) for DGR-variable proteins, and red shaded box indicates non-coding DGR template region (TR). **(B)** Box plot illustrating the expression levels of the DGR template region (DGR_TR) across the 12 timepoints (light = orange, dark = purple). **(C)** Nucleotide diversity (calculated as pi diversity) within Variable Region 1 (VR1) compared to the template repeat (TR). Adenine positions are indicated by pink shading. **(D)** Nucleotide diversity (calculated as pi diversity) within Variable Region 2 (VR2) compared to the template repeat (TR). Adenine positions are indicated by pink shading.

Strikingly, TR-RNA appears to be very highly expressed (**Figure 5B**), which is consistent with past observations (46, 47), yet unexpected for a domesticated model organism. This RNA serves a dual function – up and downstream regions are predicted to form part of the ribonucleoprotein structure of a DGR complex (48), whereas the core TR sequence contains coding information transmitted to the variable target(s) via reverse transcription and retrohoming. In a previous study with the marine filamentous cyanobacterium, *Trichodesmium erythraeum*, multiple DGR loci were identified in which TR-RNA appeared highly expressed (46). However, adenine mutation was not detected, which was attributed to long term culture stability and loss of selective pressure for targeted diversification. By contrast, adenine mutation was observed in environmental metagenomes recruited to the *Trichodesmium* reference genome, suggesting that DGR mutation is stimulated in the wild (49). Thus, prior to this study, it was unclear whether DGR mutagenesis occurs or can be detected in lab-maintained isolates of filamentous cyanobacteria.

In order to detect hypermutation in the DGR target loci, we carried out high coverage whole-genome resequencing of *N. punctiforme* from the Dark24 timepoint, chosen as a perturbation of the natural LD cycle. Resequencing revealed dynamic adenine mutation in both target genes – while detectable in the variable locus of DGR target 1 (Npun_F4889; **Figure 5C**), hypermutation is relatively higher in the variable locus of DGR target 2 (Npun_F4890; **Figure 5D**). Despite revealing a pattern consistent with adenine-specific mutation, Pi diversity is low in these loci – just above our limit of detection – suggesting that diversification is not elevated in the absence of specific environmental stress or stimuli. This is consistent with the expectation that under stable growth conditions, the cultures are not challenged with biological conflict (e.g., phage infection) or nutrient limitation. The low degree of targeted mutation is also consistent with low expression of the diversification machinery, RT and Avd. It is unclear if the perturbation to the LD cycle influences hypermutation, or if hypermutation is constitutive at low levels under general laboratory conditions. Future experiments designed to compare hypermutation between LD and LL cycles can address this question. These findings also raise the question on whether regulation of hypermutation is decoupled from the expression of the target genes themselves.

While little is known about the functional role of hypermutation target genes of cyanobacterial DGRs, they have been recently hypothesized to coordinate immunity or defense as part of multi-protein conflict mitigation systems (50). To determine whether co-localized genes in the DGR neighborhood were co-expressed, we conducted a global Pearson correlation analysis across pairwise sets of genes (**Supplemental Table S7**). This uncovered a group of apparently correlated genes, comprising a CHAT domain-containing protein (Npun_F4885), a MoxR ATPase (Npun_F4887), a VWA domain-containing protein (Npun_F4888), and DGR target 2 (Npun_F4890). Moreover, expression of DGR-RT (Npun_F4892) appears correlated with a downstream restriction endonuclease (Npun_F4897). It is perhaps surprising that putative Avd and DGR-RT were not correlated, however, this is likely a result of their overall low expression across the time series.

The co-occurrence of MoxR and VWA proteins (known as STAND-family ATPases) with DGR hypervariable targets was proposed to have a role in adaptive defense through diversifying antigen sensors, to mitigate conflict such as phage infection (50). Our findings offer first experimental evidence that co-localized STAND ATPase and DGR target genes are indeed co-expressed. As such, targeted hypermutation of putative conflict systems likely confers a fitness advantage in wild filamentous cyanobacteria. While the specific cellular function of these multipartite systems is somewhat elusive, DGRs could offer phenotypic heterogeneity and subpopulation stability during biological conflict.

## Concluding Remarks

This study provides a comprehensive view of the expression dynamics in *Nostoc punctiforme* during its natural diel cycle, highlighting a clear shift in day-to-night gene expression for key processes such as central carbon metabolism, nucleotide and amino acid conversion, and DNA repair. A combined workflow of clustering, network, and sequence diversity analyses revealed a dynamic mobilome consisting of transposons, prophage, and retroelements that drive hypermutation. Despite long-term domestication under continuous light and stable culture conditions, this multicellular cyanobacterium still partitions its gene expression into light– and dark-associated processes. Co-occurring sugar metabolism and peptidoglycan biosynthesis, along with genes for hormogonium development and motility, are preferentially expressed during light conditions. Conversely, dark conditions result in preferential expression of chromosome repair and nucleotide metabolism pathways. Throughout this temporal study, several mobile genetic elements, including transposons and putative prophage, demonstrated diel-variable expression. Additionally, we confirmed the dynamic expression diversity-generating retroelements, accompanied by adenine-specific hypermutation in target genes. Collectively, these findings offer significant insights into the regulatory complexity and genotypic plasticity of this model organism.

Some limitations present when contextualizing the transcriptome of *N. punctiforme* – especially where detailed information on diel regulation is available for unicellular cyanobacteria that may not apply in multicellular counterparts. For example, while *Synechococcus* spp. exhibit markedly lower levels of mRNA at night (38, 51), *Anabaena* spp. maintain constant mRNA concentrations throughout light-dark cycles (35), exemplifying key differences in expression between unicellular and multicellular cyanobacteria. Furthermore, current knowledge of circadian processes is predominantly described at the translational level, as the core clock mechanism is proteinaceous. Nevertheless, the circadian clock is regulated by transcriptional factors that respond to diel shifts in cellular metabolism. Instances of delayed mRNA circadian signals, likely due to post-transcriptional regulation (34), further complicate interpretation of expression data. As such, future efforts will benefit from the deconvolution of multicellular circadian and diel processes, both in terms of transcription and protein (e.g., phosphorylation) states. Overall, this work provides new insights into the natural division of cellular processes in a model cyanobacterium between day and night, and we anticipate that these findings will contribute to a deeper understanding of the regulatory complexity in multicellular cyanobacteria.

## Materials and Methods

### Bacterial culture conditions

*Nostoc punctiforme* PCC 73102 (ATCC 29133) was grown at 28°C under constant shaking conditions (130 rpm) within Allan and Arnon standard growth media, diluted fourfold (AA/4) (52). Cultures were grown under constant light (40 – 50 μmol photons m^-2^s^-1^) for two weeks to accumulate biomass prior to partitioning into replicate cultures equivalent to ∼47 μg chlorophyll. Diurnal entrainment was simulated via a 12 hour Light / 12 hour Dark (LD) growth regime, and triplicate cultures were collected per examined timepoint. To collect the final two samples (Light24 and Dark24), the 12/12 LD regime was disrupted 192 hours into the experiment (∼Pre-Dusk2). At this time, a 24 hour extended LL incubation was performed, with Dark24-designated replicate cultures covered in tin foil to occlude light (24 hour DD).

### Sample collection and nucleic acid extraction

Sample collection and nucleic acid extraction were performed as previously described (25, 53). Briefly, cultures were harvested via centrifugation and bacterial pellet cryopreservation at –80°C. Frozen pellets were subjected to lysis via bead-beating, and total RNA was precipitated with lithium chloride precipitation. RNA was further purified with the QIAGEN RNeasy Plus Mini Kit and on-column DNase treatment (QIAGEN RNase-Free DNase Set), per manufacturer’s instructions.

Genomic DNA was similarly extracted from frozen bacterial pellets. Pellets were subjected to initial lysis with lysozyme and bead-beating; total DNA was isolated following Zymo Research Quick-DNA Miniprep Plus Kit manufacturer instructions.

### Transcriptome Sequencing

Enrichment of bacterial mRNA was performed using the NEBNext rRNA Depletion Kit (Bacteria) and RNA Sample Purification Beads. Non-directional cDNA libraries were synthesized using the Invitrogen SuperScript IV First-Strand Synthesis System. Library preparation and sequencing were performed by the Keck Sequencing Facility at the Marine Biological Laboratory. Briefly, sequencing libraries were prepared using the Tecan Life Sciences Ovation® Ultralow V2 DNA-Seq Library Preparation Kit, followed by two Illumina NextSeq 500 sequencing runs (**Figure 1A**: Stages I and II; Stages III and IV). Library amplification was carried out with 10 to 14 cycles (10 cycles for replicate i, and 14 cycles for replicates ii and iii across all timepoints). Single-end reads were generated using the Illumina NextSeq 500/550 High Output Kit v2.5 (150 cycles).

### Whole genome sequencing

One biological replicate of genomic DNA extracted from the Dark24 timepoint was selected for whole genome sequencing. Library preparation was conducted as above (i.e. for RNA seq libraries), but with 7 cycles of library amplification. Sequencing was performed by the Keck Sequencing Facility at the Marine Biological Laboratory library. Paired-end reads were generated using the Illumina NextSeq 500/550 High Output Kit v2.5 (300 cycles).

### Bioinformatics

Rockhopper was used with default parameters for the alignment, assembly, and normalization of sequencing data (54)-(55). An average of 19,993,756 reads were mapped to the *N. punctiforme* genome (NCBI RefSeq assembly <u>GCF_000020025.1</u>) for all replicates and timepoints.

Overall visualizations of RNAseq data (Spearman’s correlations and principal component analysis) were calculated within R for upper quartile normalized gene counts of all samples. The bioinformatic workflow is presented within **Supplemental Figure S3**, wherein the global transcriptome was first filtered to remove low expression (i.e. <10 expression in all timepoints). Parsed expression values were Z-score normalized per gene before hierarchical clustering (distances: Euclidean; linkaging: complete; R package ComplexHeatmap v2.16.0 (56, 57)).

The weighted gene co-expressed gene networks (WGCNA) derived from all 12 experimental transcriptomes were generated with R package WGCNA v1.72-5 (58, 59). Pearson correlation values were calculated to examine co-expressed gene networks (“modules”) correlations with timepoints. Cytoscape v3.9.1 (60) was used to visualize modules, and cytoHubba v0.1 (61) was used to calculate maximal clique centrality (MCC). Broadly, “day” encompassed Pre-Dusk, MidLight, and Light24 timepoints, whereas “night” referred to Pre-Dawn, MidDark, and Dark24 timepoints.

We examined the “core” diurnal (LD) entrainment transcriptomes (**Figure 1A**, Stages II and III) for diel-variable expression. Genes having consistently low expression were filtered (as previously), prior to an analysis of variance (ANOVA) of the four states (MidDark, Pre-Dawn, MidLight, Pre-Dusk) in duplicate for the core eight transcriptomes, with Benjamini-Hochberg (BH) correction. Hierarchical clustering of significant genes (BH-adjusted p<0.05) revealed distinct “day” (Pre-Dusk and MidLight) and “night” (Pre-Dawn and MidDark) associated partitioning into two major clusters. These “day” and “night” associated genes, in combination with the previously identified WGCNA modules most-correlated with each timepoint, were submitted for gene set enrichment analysis (GSEA; R package clusterProfiler v4.8.3 ((62, 63)) and *<u>N. punctiforme</u>* <u>KEGG entry</u>) to identify “day” and “night” associated cellular processes.

Reciprocal BLASTn (v2.12.0+; default settings) analyses of clock-controlled *Nostoc* sp. PCC 7120 (NCBI RefSeq assembly <u>GCF_000009705.1</u>) genes (35) and *N. punctiforme* coding sequences were performed to identify homologues, with potential cyclical expression visualized within KEGG pathway-unique polar plots.

*Synechococcus elongatus* PCC 7942 CikA (Synpcc7942_0644; UniProt entry <u>Q9KHI5</u>) GAF region (residues 184-338) and histidine kinase domain (residues 390-611) were separately submitted to two BLASTp searches restricted to NCBI-available cyanobacteriota (v2.12.0+; Taxonomy ID 1117; default settings). Clustal Omega (v1.2.3; default settings) was used to align proteins containing both domains, generating a consensus sequence that was used for a tBLASTn (v2.12.0+; default settings) search of *N. punctiforme* genome, with a 50% query coverage threshold for determining CikA-like histidine kinases.

Whole genomic sequencing reads were aligned to the *N. punctiforme* genome (NCBI RefSeq assembly <u>GCF_000020025.1</u>) using bwa-mem2 (64). Mapping file sorting and coverage were completed with Samtools v1.17 (65). A custom python script was developed to calculate the pi diversity of the predicted DGR hypermutable region.

### RT-qPCR

A separate diel experiment was performed for reverse transcription-quantitative PCR (RT-qPCR) examination of *N. punctiforme* expression. Eighteen replicate flasks of AA/4 were inoculated with biomass equivalent to ∼51 μg chlorophyll before incubation at repeated growth conditions (12 L/12 D cycle, light intensity, temperature, shaking conditions). Cultures were grown for 47.5 hours prior to sample collection (as previously described) for the Pre-Dawn timepoint. Subsequent samples were collected every 6.5 hours for the MidLight, Pre-Dusk, and Mid-Dark timepoints, yielding triplicate replicates per timepoint. To collect the final two samples (Light24 and Dark24), the 12/12 LD regime was disrupted 67 hours into the experiment (∼MidDark). At this time, a 24 hour extended LL incubation was performed, with Dark24 designated replicate cultures covered in tin foil to occlude light (24 hour DD).

RT-qPCR samples were collected and processed similarly to the RNAseq workflow. The Applied Biosystems QuantStudio 5 Real-Time PCR System and the NEB Luna Universal qPCR Master Mix were used to quantify selected genes’ expressions. In addition to the repeated diel growth samples (n=18), the last 18 RNAseq samples were submitted for RT-qPCR (**Figure 1A**; Stages III and IV). Normalization was performed with *rnpB* (Npun_r018; RNA component of RNaseP; F 5’-TAAGAGCGCACCAGCAGTAT – 3’ and R 5’-

CATTGAGCGGAACTGGTAAA – 3’). Chosen experimental genes were *murA* (Npun_R5719; UDP-N-acetylglucosamine 1-carboxyvinyltransferase; F 5’-CCTCGCACATGCAAGTCAAC – 3’ and R 5’ – TCGCCATTGGAGCCATTCTT – 3’); *psbF* (Npun_F5552; photosystem II cytochrome b559 subunit beta; F 5’-AGCGGCAATAACATCAATCAACC – 3’ and R 5’ – AAATTGCATTGCGGCGATCG – 3’); and a phage tail sheath protein gene (Npun_F1112; F 5’-TACTTTGCCCGTCCCAACTC – 3’ and R 5’-AACGACCGCCACCATTCATA – 3’). Standard curve and absolute quantification calculations were performed within Applied Biosystems Design & Analysis Software v2.7.0.

### Data availability

Transcriptome raw data, metadata, and gene counts were uploaded to the GEO database with the accession numbers XX_YY, WGSeq raw reads were submitted to the SRA database and have accession numbers YY_ZZ. R and python scripts are available on GitHub.

## Acknowledgements

We thank Bailey Fallon, Callie Herschfield, and Hilary Morrison, and the Keck Sequencing Facility at the Marine Biological Laboratory for generating sequence libraries. Computational resources were provided by the Bay Paul Center for Comparative Molecular Biology and Evolution (MBL). This work was supported by the G. Unger Vetlesen Foundation and the Owens Family Foundation.

## References

1. B. E. Schirrmeister, A. Antonelli, H. C. Bagheri, The origin of multicellularity in cyanobacteria. BMC Evol. Biol. 11, 45 (2011).

2. J. Lehner, et al., The morphogene AmiC2 is pivotal for multicellular development in the cyanobacterium Nostoc punctiforme. Mol. Microbiol. 79, 1655–1669 (2011).

3. J. C. Meeks, E. L. Campbell, M. L. Summers, F. C. Wong, Cellular differentiation in the cyanobacterium Nostoc punctiforme. Arch. Microbiol. 178, 395–403 (2002).

4. D. D. Risser, Hormogonium Development and Motility in Filamentous Cyanobacteria. Appl. Environ. Microbiol. 89 (2023).

5. C. Argueta, M. L. Summers, Characterization of a model system for the study of Nostoc punctiforme akinetes. Arch. Microbiol. 183, 338–346 (2005).

6. S. Belton, P. F. McCabe, C. K. Y. Ng, The cyanobacterium, Nostoc punctiforme can protect against programmed cell death and induce defence genes in Arabidopsis thaliana. J. Plant Interact. 16, 64–74 (2021).

7. L. E. Moraes, et al., Resequencing and annotation of the Nostoc punctiforme ATTC 29133 genome: facilitating biofuel and high-value chemical production. AMB Express 7, 42 (2017).

8. R. Garg, I. Maldener, The Formation of Spore-Like Akinetes: A Survival Strategy of Filamentous Cyanobacteria. Microb Physiol 31, 296–305 (2021).

9. K. Klicki, D. Ferreira, D. Risser, F. Garcia-Pichel, A regulatory linkage between scytonemin production and hormogonia differentiation in. iScience 25, 104361 (2022).

10. A. Herrero, J. Stavans, E. Flores, The multicellular nature of filamentous heterocyst-forming cyanobacteria. FEMS Microbiol. Rev. 40, 831–854 (2016).

11. J. Toepel, E. Welsh, T. C. Summerfield, H. B. Pakrasi, L. A. Sherman, Differential transcriptional analysis of the cyanobacterium Cyanothece sp. strain ATCC 51142 during light-dark and continuous-light growth. J. Bacteriol. 190, 3904–3913 (2008).

12. D. Mella-Flores, et al., Prochlorococcus and Synechococcus have Evolved Different Adaptive Mechanisms to Cope with Light and UV Stress. Front. Microbiol. 3, 285 (2012).

13. T. Elvitigala, J. Stöckel, B. K. Ghosh, H. B. Pakrasi, Effect of continuous light on diurnal rhythms in Cyanothece sp. ATCC 51142. BMC Genomics 10, 226 (2009).

14. C. H. Johnson, M. J. Rust, Circadian Rhythms in Bacteria and Microbiomes (Springer Nature, 2021).

15. V. Vijayan, R. Zuzow, E. K. O’Shea, Oscillations in supercoiling drive circadian gene expression in cyanobacteria. Proc. Natl. Acad. Sci. U. S. A. 106, 22564–22568 (2009).

16. X. Zhang, et al., Genome-wide survey of putative serine/threonine protein kinases in cyanobacteria. BMC Genomics 8, 395 (2007).

17. A. Vallota-Eastman, et al., Role of diversity-generating retroelements for regulatory pathway tuning in cyanobacteria. BMC Genomics 21, 664 (2020).

18. A. I. Garber, E. B. Sano, A. L. Gallagher, S. R. Miller, Duplicate Gene Expression and Possible Mechanisms of Paralog Retention During Bacterial Genome Expansion. Genome Biol. Evol. 16 (2024).

19. Role of the GlgX protein in glycogen metabolism of the cyanobacterium, Synechococcus elongatus PCC 7942. Biochimica et Biophysica Acta (BBA) – General Subjects 1770, 763–773 (2007).

20. M. Gründel, R. Scheunemann, W. Lockau, Y. Zilliges, Impaired glycogen synthesis causes metabolic overflow reactions and affects stress responses in the cyanobacterium Synechocystis sp. PCC 6803. Microbiology 158, 3032–3043 (2012).

21. G. Dong, et al., Elevated ATPase activity of KaiC applies a circadian checkpoint on cell division in Synechococcus elongatus. Cell 140, 529–539 (2010).

22. Dynamic Inventory of Intermediate Metabolites of Cyanobacteria in a Diurnal Cycle. iScience 23, 101704 (2020).

23. D. G. Welkie, et al., Genome-wide fitness assessment during diurnal growth reveals an expanded role of the cyanobacterial circadian clock protein KaiA. Proc. Natl. Acad. Sci. U. S. A. 115, E7174–E7183 (2018).

24. E. L. Campbell, et al., Genetic Analysis Reveals the Identity of the Photoreceptor for Phototaxis in Hormogonium Filaments of Nostoc punctiforme. J. Bacteriol. (2014). 10.1128/jb.02374-14.

25. A. Gonzalez, K. W. Riley, T. V. Harwood, E. G. Zuniga, D. D. Risser, A Tripartite, Hierarchical Sigma Factor Cascade Promotes Hormogonium Development in the Filamentous Cyanobacterium Nostoc punctiforme. mSphere 4 (2019).

26. A. M. Puszynska, E. K. O’Shea, Switching of metabolic programs in response to light availability is an essential function of the cyanobacterial circadian output pathway. Elife 6 (2017).

27. M. Hanai, et al., The Effects of Dark Incubation on Cellular Metabolism of the Wild Type Cyanobacterium Synechocystis sp. PCC 6803 and a Mutant Lacking the Transcriptional Regulator cyAbrB2. Life 4, 770–787 (2014).

28. S. S. Golden, L. A. Sherman, Optimal conditions for genetic transformation of the cyanobacterium Anacystis nidulans R2. J. Bacteriol. 158, 36–42 (1984).

29. A. Taton, et al., The circadian clock and darkness control natural competence in cyanobacteria. Nat. Commun. 11, 1688 (2020).

30. R. G. Lloyd, C. J. Rudolph, 25 years on and no end in sight: a perspective on the role of RecG protein. Curr. Genet. 62, 827–840 (2016).

31. C. Cassier-Chauvat, T. Veaudor, F. Chauvat, Comparative Genomics of DNA Recombination and Repair in Cyanobacteria: Biotechnological Implications. Front. Microbiol. 7, 1809 (2016).

32. D. Jeruzalmi, M. O’Donnell, J. Kuriyan, Crystal structure of the processivity clamp loader gamma (gamma) complex of E. coli DNA polymerase III. Cell 106, 429–441 (2001).

33. R. T. Pomerantz, M. O’Donnell, Replisome mechanics: insights into a twin DNA polymerase machine. Trends Microbiol. 15, 156–164 (2007).

34. A. C. L. Guerreiro, et al., Daily rhythms in the cyanobacterium synechococcus elongatus probed by high-resolution mass spectrometry-based proteomics reveals a small defined set of cyclic proteins. Mol. Cell. Proteomics 13, 2042–2055 (2014).

35. H. Kushige, et al., Genome-wide and heterocyst-specific circadian gene expression in the filamentous Cyanobacterium Anabaena sp. strain PCC 7120. J. Bacteriol. 195, 1276–1284 (2013).

36. R. Arbel-Goren, et al., Robust, coherent, and synchronized circadian clock-controlled oscillations along Anabaena filaments. (2021). 10.7554/eLife.64348.

37. J. C. Meeks, et al., An overview of the genome of Nostoc punctiforme, a multicellular, symbiotic cyanobacterium. Photosynth. Res. 70, 85–106 (2001).

38. H. Ito, et al., Cyanobacterial daily life with Kai-based circadian and diurnal genome-wide transcriptional control in Synechococcus elongatus. Proc. Natl. Acad. Sci. U. S. A. 106, 14168–14173 (2009).

39. R. P. Novick, G. E. Christie, J. R. Penadés, The phage-related chromosomal islands of Gram-positive bacteria. Nat. Rev. Microbiol. 8, 541–551 (2010).

40. M. Cervera-Alamar, et al., Mobilisation Mechanism of Pathogenicity Islands by Endogenous Phages in Staphylococcus aureus clinical strains. Sci. Rep. 8, 16742 (2018).

41. B. R. Macadangdang, S. K. Makanani, J. F. Miller, Accelerated Evolution by Diversity-Generating Retroelements. Annu. Rev. Microbiol. 76, 389–411 (2022).

42. T. Hackl, et al., Novel integrative elements and genomic plasticity in ocean ecosystems. Cell 186, 47–62.e16 (2023).

43. A. C. Baker, et al., Identification of a diagnostic marker to detect freshwater cyanophages of filamentous cyanobacteria. Appl. Environ. Microbiol. 72, 5713–5719 (2006).

44. N. T. Hu, T. Thiel, T. H. Giddings Jr, C. P. Wolk, New Anabaena and Nostoc cyanophages from sewage settling ponds. Virology 114, 236–246 (1981).

45. C. Chénard, J. F. Wirth, C. A. Suttle, Viruses Infecting a Freshwater Filamentous Cyanobacterium (Nostoc sp.) Encode a Functional CRISPR Array and a Proteobacterial DNA Polymerase B. MBio 7 (2016).

46. U. Pfreundt, M. Kopf, N. Belkin, I. Berman-Frank, W. R. Hess, The primary transcriptome of the marine diazotroph Trichodesmium erythraeum IMS101. Sci. Rep. 4, 6187 (2014).

47. B. G. Paul, et al., Retroelement-guided protein diversification abounds in vast lineages of Bacteria and Archaea. Nat Microbiol 2, 17045 (2017).

48. S. Handa, T. Biswas, J. Chakraborty, B. G. Paul, P. Ghosh, Structural Requirements for Reverse Transcription by a Diversity-generating Retroelement. bioRxiv (2023). 10.1101/2023.10.23.563531.

49. B. G. Paul, A. M. Eren, Eco-evolutionary significance of domesticated retroelements in microbial genomes. Mob. DNA 13, 6 (2022).

50. H. Doré, et al., Targeted hypermutation of putative antigen sensors in multicellular bacteria. Proc. Natl. Acad. Sci. U. S. A. 121, e2316469121 (2024).

51. S. Takano, J. Tomita, K. Sonoike, H. Iwasaki, The initiation of nocturnal dormancy in Synechococcus as an active process. BMC Biol. 13, 36 (2015).

52. C. S. Enderlin, J. C. Meeks, Pure culture and reconstitution of the Anthoceros-Nostoc symbiotic association. Planta 158, 157–165 (1983).

53. E. L. Campbell, M. L. Summers, H. Christman, M. E. Martin, J. C. Meeks, Global gene expression patterns of Nostoc punctiforme in steady-state dinitrogen-grown heterocyst-containing cultures and at single time points during the differentiation of akinetes and hormogonia. J. Bacteriol. 189, 5247–5256 (2007).

54. A computational system for identifying operons based on RNA-seq data. Methods 176, 62–70 (2020).

55. R. McClure, et al., Computational analysis of bacterial RNA-Seq data. Nucleic Acids Res. 41, e140–e140 (2013).

56. Z. Gu, R. Eils, M. Schlesner, Complex heatmaps reveal patterns and correlations in multidimensional genomic data. Bioinformatics 32, 2847–2849 (2016).

57. Z. Gu, Complex heatmap visualization. Imeta 1 (2022).

58. B. Zhang, S. Horvath, A general framework for weighted gene co-expression network analysis. Stat. Appl. Genet. Mol. Biol. 4, Article17 (2005).

59. P. Langfelder, S. Horvath, WGCNA: an R package for weighted correlation network analysis. BMC Bioinformatics 9, 1–13 (2008).

60. P. Shannon, et al., Cytoscape: a software environment for integrated models of biomolecular interaction networks. Genome Res. 13, 2498–2504 (2003).

61. C.-H. Chin, et al., cytoHubba: identifying hub objects and sub-networks from complex interactome. BMC Syst. Biol. 8 **Suppl 4**, S11 (2014).

62. clusterProfiler 4.0: A universal enrichment tool for interpreting omics data. Innov. J. 2, 100141 (2021).

63. G. Yu, L.-G. Wang, Y. Han, Q.-Y. He, clusterProfiler: an R Package for Comparing Biological Themes Among Gene Clusters. (2012). 10.1089/omi.2011.0118.

64. M. Vasimuddin, S. Misra, H. Li, S. Aluru, Efficient architecture-aware acceleration of BWA-MEM for multicore systems in 2019 IEEE International Parallel and Distributed Processing Symposium (IPDPS), (IEEE, 2019).

65. P. Danecek, et al., Twelve years of SAMtools and BCFtools. Gigascience 10 (2021).

