## Supplemental Table S1 for "Diel expression dynamics in filamentous cyanobacteria"

**Supplemental Table S1.** Correlations of sugar metabolism and peptidoglycan biosynthesis gene expressions.

| Gene1 | Gene2 | CorrelationValue | ComparedGenes |
|---|---|---|---|
| Npun_R2486 | Npun_R5719 | 0.963800567 | *murE-murA* |
| Npun_R1733 | Npun_R1952 | 0.949753362 | *LMW PBP-uppS* |
| Npun_F0446 | Npun_F0447 | 0.945244621 | *murC-murB* |
| Npun_F3659 | Npun_F5597 | 0.911178829 | ***rpaA*** *-ddl* |
| Npun_F4452 | Npun_R3281 | 0.876634722 | Class B PBP-Class A PBP |
| Npun_R1302 | Npun_R1733 | 0.866959791 | ***nagB*** - LMW PBP |
| Npun_F2411 | Npun_R4507 | 0.858599638 | *murG-bacA* |
| Npun_F0907 | Npun_R5719 | 0.857098146 | ***glmU-*** *murA* |
| Npun_F0907 | Npun_R4507 | 0.833793629 | ***glmU*** *-bacA* |
| Npun_F0907 | Npun_R2557 | 0.833598268 | ***glmU*** -LMW PBP |
| Npun_F4453 | Npun_R5719 | 0.829890155 | Class B PBP-*murA* |
| Npun_F4453 | Npun_R4507 | 0.822625392 | Class B PBP-*bacA* |
| Npun_F0168 | Npun_R3559 | 0.821610231 | Class B PBP-*murT* |
| Npun_F0168 | Npun_R2557 | 0.815816275 | Class B PBP-LMW PBP |
| Npun_F0447 | Npun_R3559 | 0.799008701 | *murB-murT* |
| Npun_R2557 | Npun_R5719 | 0.798527088 | LMW PBP-*murA* |
| Npun_R2557 | Npun_R4507 | 0.792949227 | LMW PBP-*bacA* |
| Npun_F0907 | Npun_F4453 | 0.786277386 | ***glmU*** -Class B PBP |
| Npun_F0446 | Npun_R3559 | 0.779678986 | *murC-murT* |
| Npun_F0168 | Npun_F0907 | 0.774839495 | Class B PBP-***glmU*** |
| Npun_F2411 | Npun_F4453 | 0.77106193 | *murG*- Class B PBP |
| Npun_F2411 | Npun_R5719 | 0.764573276 | *murG-murA* |
| Npun_R1302 | Npun_R1952 | 0.760076043 | ***nagB*** *-uppS* |
| Npun_F4453 | Npun_R2486 | 0.758140407 | Class B PBP-*murE* |
| Npun_F3925 | Npun_F3940 | 0.750430543 | *pgi-mraY* |

**Non-peptidoglycan Gene**
