## Supplemental Table S2 for "Diel expression dynamics in filamentous cyanobacteria"

**Supplemental Table S2.** Correlations of cell division, peptidoglycan biosynthesis, and sugar metabolism gene expressions.

| Gene1 | Gene2 | CorrelationValue | ComparedGenes |
|---|---|---|---|
| Npun_R2486 | Npun_R5719 | 0.963800567 | MurE-MurA |
| Npun_F5138 | Npun_R1839 | 0.954991591 | FtsE-MreD |
| Npun_R1733 | Npun_R1952 | 0.949753362 | LMW PBP-UppS |
| Npun_F3647 | Npun_F3659 | 0.946919803 | MinC-RpaA |
| Npun_F0446 | Npun_F0447 | 0.945244621 | MurC-MurB |
| Npun_F5597 | Npun_R4092 | 0.938576687 | Ddl-FtsK |
| Npun_F4453 | Npun_R4933 | 0.929808899 | Class B PBP-Cdv1 |
| Npun_F3647 | Npun_R0056 | 0.921719228 | MinC-DUF152 |
| Npun_R1698 | Npun_R2486 | 0.921037611 | SepF-MurE |
| Npun_F3659 | Npun_F5597 | 0.911178829 | RpaA-Ddl |
| Npun_BF043 | Npun_F0907 | 0.906259505 | MinD-GlmU |
| Npun_F5138 | Npun_R1733 | 0.900532737 | FtsE-LMW PBP |
| Npun_F4452 | Npun_R1841 | 0.898245014 | Class B PBP-MreB |
| Npun_R2486 | Npun_R4806 | 0.897717901 | MurE-FtsQ |
| Npun_F4881 | Npun_F5597 | 0.897189533 | FtsH-Ddl |
| Npun_F3647 | Npun_F5597 | 0.894821592 | MinC-Ddl |
| Npun_R1698 | Npun_R4806 | 0.89448759 | SepF-FtsQ |
| Npun_F3659 | Npun_R4092 | 0.891497607 | RpaA-FtsK |
| Npun_F5214 | Npun_R0056 | 0.889627672 | GlmS-DUF152 |
| Npun_R4806 | Npun_R5719 | 0.889400819 | FtsQ-MurA |
| Npun_F5138 | Npun_R1952 | 0.886711221 | FtsE-UppS |
| Npun_F3659 | Npun_R6629 | 0.881969897 | RpaA-SepI |
| Npun_R1698 | Npun_R4804 | 0.880481335 | SepF-FtsZ |
| Npun_R1840 | Npun_R1841 | 0.879328505 | MreC-MreB |
| Npun_R1698 | Npun_R5719 | 0.878051425 | SepF-MurA |
| Npun_F4452 | Npun_R3281 | 0.876634722 | Class B PBP-Class A PBP |
| Npun_F4881 | Npun_R4092 | 0.875996454 | FtsH-FtsK |
| Npun_R4933 | Npun_R5149 | 0.875761242 | Cdv1-GAF-HisKin |
| Npun_R4507 | Npun_R5149 | 0.875678492 | BacA-GAF-HisKin |
| Npun_F2411 | Npun_R5149 | 0.875162279 | MurG-GAF-HisKin |
| Npun_R6354 | Npun_R6629 | 0.874779885 | YlmH-SepI |
| Npun_R1733 | Npun_R1839 | 0.870970857 | LMW PBP-MreD |
| Npun_R1302 | Npun_R1733 | 0.866959791 | NagB-LMW PBP |
| Npun_R1839 | Npun_R1841 | 0.866074827 | MreD-MreB |
| Npun_R4507 | Npun_R4933 | 0.863726036 | BacA-Cdv1 |
| Npun_F3659 | Npun_R0056 | 0.863131187 | RpaA-DUF152 |
| Npun_BF043 | Npun_R2557 | 0.860702956 | MinD-LMW PBP |
| Npun_F3648 | Npun_F3649 | 0.860169981 | MinD-MinE |
| Npun_F3647 | Npun_R6629 | 0.859454021 | MinC-SepI |
| Npun_F2411 | Npun_R4507 | 0.858599638 | MurG-BacA |
| Npun_BF043 | Npun_F0168 | 0.858532054 | MinD-Class B PBP |
| Npun_R1839 | Npun_R1840 | 0.857190145 | MreD-MreC |
| Npun_F0907 | Npun_R5719 | 0.857098146 | GlmU-MurA |
| Npun_R1733 | Npun_R1841 | 0.853753075 | LMW PBP-MreB |
| Npun_F3647 | Npun_R4092 | 0.845587208 | MinC-FtsK |
| Npun_R1698 | Npun_R1841 | 0.84248621 | SepF-MreB |
| Npun_F2411 | Npun_R4933 | 0.840462196 | MurG-Cdv1 |
| Npun_F3659 | Npun_R6354 | 0.838489013 | RpaA-YlmH |
| Npun_R1839 | Npun_R1952 | 0.838013328 | MreD-UppS |
| Npun_F5597 | Npun_R0056 | 0.835556776 | Ddl-DUF152 |
| Npun_F0907 | Npun_R4507 | 0.833793629 | GlmU-BacA |
| Npun_F0907 | Npun_R2557 | 0.833598268 | GlmU-LMW PBP |
| Npun_F5597 | Npun_R6354 | 0.832505749 | Ddl-YlmH |
| Npun_F4453 | Npun_R5719 | 0.829890155 | Class B PBP-MurA |
| Npun_F0907 | Npun_R5149 | 0.82564693 | GlmU-GAF-HisKin |
| Npun_F4453 | Npun_R4507 | 0.822625392 | Class B PBP-BacA |
| Npun_F0168 | Npun_R3559 | 0.821610231 | Class B PBP-MurT |
| Npun_R4092 | Npun_R6354 | 0.821325661 | FtsK-YlmH |
| Npun_R1841 | Npun_R1952 | 0.820090948 | MreB-UppS |
| Npun_F0447 | Npun_R4806 | 0.819999098 | MurB-FtsQ |
| Npun_F0168 | Npun_R2557 | 0.815816275 | Class B PBP-LMW PBP |
| Npun_F5138 | Npun_R1840 | 0.807280548 | FtsE-MreC |
| Npun_F3659 | Npun_F4881 | 0.805513371 | RpaA-FtsH |
| Npun_F0907 | Npun_R4806 | 0.805274401 | GlmU-FtsQ |
| Npun_F5138 | Npun_R1841 | 0.803919879 | FtsE-MreB |
| Npun_F0447 | Npun_R3559 | 0.799008701 | MurB-MurT |
| Npun_R2557 | Npun_R5719 | 0.798527088 | LMW PBP-MurA |
| Npun_R2486 | Npun_R4804 | 0.797299888 | MurE-FtsZ |
| Npun_F0907 | Npun_R4804 | 0.796054301 | GlmU-FtsZ |
| Npun_R1698 | Npun_R1840 | 0.79543787 | SepF-MreC |
| Npun_R2557 | Npun_R4507 | 0.792949227 | LMW PBP-BacA |
| Npun_R4804 | Npun_R5719 | 0.792599137 | FtsZ-MurA |
| Npun_F3647 | Npun_R6354 | 0.792508293 | MinC-YlmH |
| Npun_F5597 | Npun_R6629 | 0.789144362 | Ddl-SepI |
| Npun_F4453 | Npun_R4804 | 0.786955439 | Class B PBP-FtsZ |
| Npun_R2557 | Npun_R4804 | 0.786854897 | LMW PBP-FtsZ |
| Npun_R1841 | Npun_R3281 | 0.786346127 | MreB-Class A PBP |
| Npun_F0907 | Npun_F4453 | 0.786277386 | GlmU-Class B PBP |
| Npun_BF043 | Npun_R4806 | 0.785448443 | MinD-FtsQ |
| Npun_R1952 | Npun_R4806 | 0.782383771 | UppS-FtsQ |
| Npun_R1841 | Npun_R4806 | 0.78114049 | MreB-FtsQ |
| Npun_R3910 | Npun_R4092 | 0.779751668 | BolA-FtsK |
| Npun_F0446 | Npun_R3559 | 0.779678986 | MurC-MurT |
| Npun_R4933 | Npun_R5719 | 0.775928629 | Cdv1-MurA |
| Npun_F0168 | Npun_F0907 | 0.774839495 | Class B PBP-GlmU |
| Npun_F4453 | Npun_R5149 | 0.773071614 | Class B PBP-GAF-HisKin |
| Npun_F0907 | Npun_R4933 | 0.771274521 | GlmU-Cdv1 |
| Npun_F2411 | Npun_F4453 | 0.77106193 | MurG-Class B PBP |
| Npun_R4092 | Npun_R6629 | 0.770623908 | FtsK-SepI |
| Npun_R1698 | Npun_R1839 | 0.767269352 | SepF-MreD |
| Npun_BF043 | Npun_R5719 | 0.765534647 | MinD-MurA |
| Npun_F2411 | Npun_R5719 | 0.764573276 | MurG-MurA |
| Npun_F5138 | Npun_R1302 | 0.762791289 | FtsE-NagB |
| Npun_R1302 | Npun_R1952 | 0.760076043 | NagB-UppS |
| Npun_F3647 | Npun_F5214 | 0.759735932 | MinC-GlmS |
| Npun_F3647 | Npun_F4881 | 0.758607248 | MinC-FtsH |
| Npun_R4804 | Npun_R4806 | 0.758602916 | FtsZ-FtsQ |
| Npun_F4453 | Npun_R2486 | 0.758140407 | Class B PBP-MurE |
| Npun_BF043 | Npun_R4009 | 0.75431246 | MinD-MurD |
| Npun_F4452 | Npun_R1839 | 0.752967292 | Class B PBP-MreD |
| Npun_F4881 | Npun_R0056 | 0.752668922 | FtsH-DUF152 |
| Npun_BF043 | Npun_R4804 | 0.751019748 | MinD-FtsZ |
| Npun_F3925 | Npun_F3940 | 0.750430543 | Pgi-MraY |
