## Supplemental Table S3 for "Diel expression dynamics in filamentous cyanobacteria"

**Supplemental Table S3.** BLASTp results of *Synechococcus elongatus* CikA query.

| Locus Tag | Accession | Name | Bit-Score | E Value | % Pairwise Identity | % Identical Sites | Hit Start | Hit End | Query Start | Query End | % Query Coverage |
|---|---|---|---|---|---|---|---|---|---|---|---|
| Npun_R1685 | ACC80356.1 | CP001037 - multi-sensor hybrid histidine kinase | 219.935 | 1.63E-60 | 32.00 | 32.00 | 2083 | 3672 | 159 | 707 | 72.81% |
| Npun_R3691 | ACC82081.1 | CP001037 - multi-sensor hybrid histidine kinase | 231.876 | 1.91E-64 | 33.70 | 33.70 | 3559 | 5163 | 159 | 682 | 69.50% |
| Npun_F5035 | ACC83378.1 | CP001037 - multi-sensor hybrid histidine kinase | 160.999 | 3.17E-41 | 28.20 | 28.20 | 3538 | 5109 | 159 | 677 | 68.83% |
| Npun_F2363 | ACC80931.1 | CP001037 - multi-sensor hybrid histidine kinase | 186.037 | 2.58E-49 | 29.80 | 29.80 | 1414 | 2943 | 159 | 626 | 62.07% |
| Npun_F1185 | ACC79910.1 | CP001037 - multi-sensor signal transduction histidine kinase | 181.03 | 2.37E-49 | 29.60 | 29.60 | 391 | 1716 | 158 | 621 | 61.54% |
| Npun_F6362 | ACC84634.1 | CP001037 - multi-sensor signal transduction histidine kinase | 155.606 | 7.85E-40 | 27.20 | 27.20 | 1249 | 2547 | 149 | 611 | 61.41% |
| Npun_F1000 | ACC79731.1 | CP001037 - GAF sensor signal transduction histidine kinase | 439.884 | 2.49E-145 | 50.50 | 50.50 | 589 | 2037 | 150 | 612 | 61.41% |
| Npun_R4776 | ACC83125.1 | CP001037 - multi-sensor signal transduction histidine kinase | 108.997 | 6.00E-25 | 23.40 | 23.40 | 1852 | 3285 | 154 | 608 | 60.34% |
| Npun_R6149 | ACC84435.1 | CP001037 - multi-sensor signal transduction histidine kinase | 133.265 | 1.13E-32 | 24.10 | 24.10 | 1210 | 2586 | 158 | 608 | 59.81% |
| Npun_R1550 | ACC80239.1 | CP001037 - GAF sensor signal transduction histidine kinase | 159.844 | 6.36E-42 | 29.80 | 29.80 | 460 | 1698 | 158 | 608 | 59.81% |
| Npun_F1203 | ACC79928.1 | CP001037 - multi-sensor signal transduction histidine kinase | 88.1965 | 1.51E-18 | 23.90 | 23.90 | 1282 | 2805 | 159 | 608 | 59.68% |
| Npun_R0896 | ACC79633.1 | CP001037 - multi-sensor hybrid histidine kinase | 161.77 | 1.25E-41 | 28.80 | 28.80 | 1312 | 2703 | 263 | 710 | 59.42% |
| Npun_F5679 | ACC83981.1 | CP001037 - multi-sensor hybrid multi-kinase | 145.206 | 4.37E-36 | 27.40 | 27.40 | 5743 | 7104 | 282 | 727 | 59.15% |
| Npun_R2903 | ACC81436.1 | CP001037 - multi-sensor signal transduction histidine kinase | 154.836 | 2.04E-39 | 26.10 | 26.10 | 2089 | 3324 | 161 | 605 | 59.02% |
| Npun_R5149 | ACC83481.1 | CP001037 - GAF sensor signal transduction histidine kinase | 153.295 | 1.66E-39 | 29.60 | 29.60 | 724 | 1959 | 165 | 608 | 58.89% |
| Npun_AF142 | ACC85008.1 | CP001038 - GAF sensor signal transduction histidine kinase | 128.642 | 4.77E-31 | 26.90 | 26.90 | 2161 | 3543 | 167 | 608 | 58.62% |
| Npun_R5113 | ACC83447.1 | CP001037 - GAF sensor signal transduction histidine kinase | 97.8265 | 1.83E-21 | 23.20 | 23.20 | 1696 | 3006 | 167 | 608 | 58.62% |
| Npun_R1597 | ACC80282.1 | CP001037 - GAF sensor signal transduction histidine kinase | 118.627 | 6.15E-28 | 25.30 | 25.30 | 1738 | 3108 | 167 | 608 | 58.62% |
| Npun_F2854 | ACC81387.1 | CP001037 - GAF sensor signal transduction histidine kinase | 129.413 | 2.66E-31 | 25.00 | 25.00 | 1741 | 3144 | 167 | 608 | 58.62% |
| Npun_R6125 | ACC84413.1 | CP001037 - multi-sensor signal transduction histidine kinase | 99.3673 | 4.80E-22 | 26.20 | 26.20 | 1060 | 2247 | 169 | 608 | 58.36% |
| Npun_R5313 | ACC83634.1 | CP001037 - multi-sensor signal transduction histidine kinase | 105.916 | 5.09E-24 | 26.00 | 26.00 | 1315 | 2697 | 170 | 608 | 58.22% |
| Npun_F2781 | ACC81314.1 | CP001037 - GAF sensor signal transduction histidine kinase | 107.842 | 9.55E-25 | 28.50 | 28.50 | 643 | 1971 | 159 | 595 | 57.96% |
| Npun_F4131 | ACC82509.1 | CP001037 - GAF sensor hybrid histidine kinase | 144.05 | 7.41E-37 | 27.50 | 27.50 | 328 | 1551 | 278 | 709 | 57.29% |
| Npun_R6464 | ACC84732.1 | CP001037 - multi-sensor hybrid histidine kinase | 161.384 | 7.12E-42 | 31.10 | 31.10 | 967 | 2118 | 293 | 698 | 53.85% |
| Npun_F5092 | ACC83426.1 | CP001037 - multi-sensor hybrid histidine kinase | 149.443 | 1.15E-37 | 27.90 | 27.90 | 2290 | 3585 | 291 | 694 | 53.58% |
| Npun_R3548 | ACC81949.1 | CP001037 - multi-sensor hybrid histidine kinase | 132.494 | 3.19E-32 | 28.30 | 28.30 | 2914 | 4230 | 302 | 694 | 52.12% |
| Npun_R5897 | ACC84193.1 | CP001037 - multi-sensor hybrid histidine kinase | 183.726 | 1.28E-48 | 30.70 | 30.70 | 3055 | 4251 | 295 | 677 | 50.80% |
| Npun_F1211 | ACC79935.1 | CP001037 - integral membrane sensor hybrid histidine kinase | 174.481 | 6.76E-46 | 34.70 | 34.70 | 676 | 1797 | 321 | 697 | 50.00% |
| Npun_R3825 | ACC82209.1 | CP001037 - hybrid histidine kinase | 211.846 | 4.84E-60 | 35.40 | 35.40 | 571 | 1698 | 331 | 706 | 49.87% |
| Npun_F1600 | ACC80285.1 | CP001037 - integral membrane sensor hybrid histidine kinase | 162.54 | 1.13E-42 | 34.40 | 34.40 | 382 | 1491 | 351 | 705 | 47.08% |
| Npun_F5479 | ACC83786.1 | CP001037 - integral membrane sensor hybrid histidine kinase | 185.267 | 1.97E-49 | 38.10 | 38.10 | 1300 | 2376 | 351 | 697 | 46.02% |
| Npun_F2686 | ACC81226.1 | CP001037 - GAF sensor hybrid histidine kinase | 184.882 | 6.58E-49 | 34.90 | 34.90 | 4510 | 5658 | 360 | 704 | 45.76% |
| Npun_R2035 | ACC80662.1 | CP001037 - PAS/PAC sensor hybrid histidine kinase | 142.51 | 1.91E-35 | 32.10 | 32.10 | 2143 | 3189 | 384 | 727 | 45.62% |
| Npun_R1432 | ACC80139.1 | CP001037 - GAF sensor signal transduction histidine kinase | 78.9518 | 4.79E-16 | 27.30 | 27.30 | 277 | 1254 | 265 | 608 | 45.62% |
| Npun_R6347 | ACC84619.1 | CP001037 - multi-sensor hybrid histidine kinase | 169.859 | 1.69E-44 | 36.50 | 36.50 | 949 | 1986 | 355 | 694 | 45.09% |
| Npun_R1798 | ACC80460.1 | CP001037 - multi-sensor hybrid histidine kinase | 169.859 | 2.89E-44 | 35.80 | 35.80 | 1876 | 2913 | 384 | 719 | 44.56% |
| Npun_R4748 | ACC83100.1 | CP001037 - multi-sensor hybrid histidine kinase | 239.58 | 3.86E-69 | 40.50 | 40.50 | 1084 | 2118 | 384 | 714 | 43.90% |
| Npun_R2262 | ACC80851.1 | CP001037 - PAS/PAC sensor hybrid histidine kinase | 134.035 | 3.23E-33 | 31.30 | 31.30 | 844 | 1869 | 384 | 710 | 43.37% |
| Npun_F6350 | ACC84622.1 | CP001037 - PAS/PAC sensor hybrid histidine kinase | 155.606 | 8.20E-40 | 34.40 | 34.40 | 1525 | 2535 | 384 | 710 | 43.37% |
| Npun_R2272 | ACC80860.1 | CP001037 - multi-sensor signal transduction multi-kinase | 100.523 | 3.67E-22 | 26.60 | 26.60 | 4387 | 5403 | 289 | 615 | 43.37% |
| Npun_F2908 | ACC81440.1 | CP001037 - multi-sensor hybrid histidine kinase | 160.614 | 3.89E-41 | 33.70 | 33.70 | 3127 | 4140 | 384 | 710 | 43.37% |
| Npun_R4028 | ACC82407.1 | CP001037 - GAF sensor signal transduction histidine kinase | 86.2705 | 2.40E-18 | 25.50 | 25.50 | 328 | 1335 | 283 | 609 | 43.37% |
| Npun_R2263 | ACC80852.1 | CP001037 - hybrid histidine kinase | 162.155 | 3.18E-42 | 34.00 | 34.00 | 649 | 1650 | 384 | 707 | 42.97% |
| Npun_R2375 | ACC80942.1 | CP001037 - multi-sensor hybrid histidine kinase | 193.741 | 5.44E-52 | 34.20 | 34.20 | 1582 | 2697 | 384 | 704 | 42.57% |
| Npun_DR038 | ACC85457.1 | CP001041 - putative PAS/PAC sensor protein | 158.688 | 1.33E-40 | 34.80 | 34.80 | 2782 | 3741 | 384 | 701 | 42.18% |
| Npun_F6040 | ACC84330.1 | CP001037 - multi-sensor signal transduction histidine kinase | 176.022 | 1.95E-46 | 34.40 | 34.40 | 1732 | 2640 | 291 | 608 | 42.18% |
| Npun_F2346 | ACC80914.1 | CP001037 - multi-sensor hybrid histidine kinase | 142.895 | 1.78E-35 | 33.00 | 33.00 | 1135 | 2016 | 311 | 626 | 41.91% |
| Npun_R1760 | ACC80422.1 | CP001037 - GAF sensor hybrid histidine kinase | 170.629 | 1.35E-45 | 35.40 | 35.40 | 661 | 1608 | 384 | 698 | 41.78% |
| Npun_R2209 | ACC80805.1 | CP001037 - GAF sensor signal transduction histidine kinase | 79.7221 | 2.46E-16 | 28.50 | 28.50 | 367 | 1269 | 296 | 608 | 41.51% |
| Npun_AR131 | ACC85002.1 | CP001038 - multi-sensor hybrid histidine kinase | 159.073 | 5.62E-41 | 36.40 | 36.40 | 1126 | 2088 | 384 | 695 | 41.38% |
| Npun_R3083 | ACC81570.1 | CP001037 - multi-sensor hybrid histidine kinase | 138.658 | 1.51E-34 | 30.80 | 30.80 | 874 | 1929 | 384 | 695 | 41.38% |
| Npun_R2268 | ACC80857.1 | CP001037 - PAS/PAC sensor hybrid histidine kinase | 146.362 | 4.36E-37 | 33.10 | 33.10 | 1033 | 1968 | 384 | 694 | 41.25% |
| Npun_R1868 | ACC80525.1 | CP001037 - multi-sensor hybrid histidine kinase | 133.65 | 5.13E-33 | 30.50 | 30.50 | 1018 | 1968 | 384 | 694 | 41.25% |
| Npun_F2346 | ACC80914.1 | CP001037 - multi-sensor hybrid histidine kinase | 143.28 | 1.24E-35 | 31.00 | 31.00 | 2611 | 3555 | 384 | 694 | 41.25% |
| Npun_R3572 | ACC81968.1 | CP001037 - multi-sensor hybrid histidine kinase | 151.369 | 3.28E-38 | 35.50 | 35.50 | 3388 | 4320 | 384 | 694 | 41.25% |
| Npun_R3591 | ACC81987.1 | CP001037 - multi-sensor hybrid histidine kinase | 164.081 | 2.21E-42 | 33.20 | 33.20 | 1987 | 2949 | 384 | 694 | 41.25% |
| Npun_F5360 | ACC83673.1 | CP001037 - multi-sensor hybrid histidine kinase | 165.236 | 1.21E-42 | 36.10 | 36.10 | 2044 | 2973 | 384 | 682 | 39.66% |
| Npun_R4744 | ACC83096.1 | CP001037 - PAS/PAC sensor hybrid histidine kinase | 150.984 | 3.62E-38 | 33.80 | 33.80 | 1948 | 2913 | 384 | 677 | 38.99% |
| Npun_F2889 | ACC81422.1 | CP001037 - CBS sensor hybrid histidine kinase | 194.512 | 3.13E-52 | 36.10 | 36.10 | 1165 | 2172 | 384 | 677 | 38.99% |
| Npun_R1449 | ACC80152.1 | CP001037 - response regulator receiver sensor signal transduction histidine kinase | 183.726 | 1.80E-51 | 38.80 | 38.80 | 337 | 1224 | 333 | 620 | 38.20% |
| Npun_R1759 | ACC80421.1 | CP001037 - GAF sensor signal transduction histidine kinase | 160.999 | 7.33E-42 | 39.00 | 39.00 | 1219 | 2058 | 322 | 608 | 38.06% |
| Npun_R3054 | ACC81549.1 | CP001037 - Chase sensor signal transduction histidine kinase | 118.242 | 7.15E-29 | 32.50 | 32.50 | 466 | 1284 | 327 | 610 | 37.67% |
| Npun_F0022 | ACC78825.1 | CP001037 - response regulator receiver sensor signal transduction histidine kinase | 128.642 | 8.95E-33 | 33.20 | 33.20 | 352 | 1149 | 335 | 608 | 36.34% |
| Npun_F1439 | ACC80146.1 | CP001037 - integral membrane sensor signal transduction histidine kinase | 150.214 | 1.87E-39 | 34.00 | 34.00 | 631 | 1413 | 342 | 608 | 35.41% |
| Npun_R1236 | ACC79960.1 | CP001037 - integral membrane sensor signal transduction histidine kinase | 143.28 | 4.92E-36 | 34.40 | 34.40 | 1033 | 1830 | 344 | 608 | 35.15% |
| Npun_R1012 | ACC79743.1 | CP001037 - integral membrane sensor signal transduction histidine kinase HepK | 177.178 | 4.82E-48 | 42.70 | 42.70 | 910 | 1707 | 350 | 608 | 34.35% |
| Npun_R3198 | ACC81651.1 | CP001037 - PAS/PAC sensor signal transduction histidine kinase | 109.383 | 2.59E-26 | 29.10 | 29.10 | 409 | 1137 | 358 | 615 | 34.22% |
| Npun_R4211 | ACC82586.1 | CP001037 - PAS/PAC sensor hybrid histidine kinase | 142.895 | 1.28E-35 | 37.10 | 37.10 | 2194 | 2982 | 384 | 641 | 34.22% |
| Npun_F6002 | ACC84292.1 | CP001037 - multi-sensor signal transduction histidine kinase | 110.153 | 6.60E-26 | 32.70 | 32.70 | 763 | 1482 | 355 | 608 | 33.69% |
| Npun_F0020 | ACC78823.1 | CP001037 - multi-sensor signal transduction histidine kinase | 96.6709 | 3.37E-21 | 30.70 | 30.70 | 1504 | 2235 | 357 | 608 | 33.42% |
| Npun_F3565 | ACC81961.1 | CP001037 - multi-sensor signal transduction multi-kinase | 103.219 | 5.58E-23 | 29.90 | 29.90 | 5428 | 6129 | 356 | 603 | 32.89% |
| Npun_F3675 | ACC82065.1 | CP001037 - multi-sensor signal transduction histidine kinase | 88.9669 | 3.73E-19 | 30.00 | 30.00 | 787 | 1479 | 359 | 603 | 32.49% |
| Npun_BF140 | ACC85272.1 | CP001039 - integral membrane sensor signal transduction histidine kinase | 99.3673 | 8.39E-23 | 30.00 | 30.00 | 574 | 1239 | 386 | 612 | 30.11% |
| Npun_R0454 | ACC79233.1 | CP001037 - GAF sensor signal transduction histidine kinase | 95.5153 | 3.49E-21 | 28.40 | 28.40 | 688 | 1518 | 384 | 610 | 30.11% |
| Npun_F0839 | ACC79577.1 | CP001037 - response regulator receiver sensor signal transduction histidine kinase | 116.701 | 1.07E-28 | 31.30 | 31.30 | 430 | 1137 | 386 | 611 | 29.97% |
| Npun_R1448 | ACC80151.1 | CP001037 - response regulator receiver sensor signal transduction histidine kinase | 117.857 | 3.33E-29 | 29.40 | 29.40 | 430 | 1119 | 386 | 610 | 29.84% |
| Npun_F0957 | ACC79688.1 | CP001037 - response regulator receiver sensor signal transduction histidine kinase | 121.324 | 2.29E-30 | 32.00 | 32.00 | 424 | 1077 | 384 | 608 | 29.84% |
| Npun_F4953 | ACC83298.1 | CP001037 - PAS/PAC sensor signal transduction histidine kinase | 129.028 | 2.13E-32 | 35.60 | 35.60 | 667 | 1326 | 384 | 608 | 29.84% |
| Npun_F3541 | ACC81945.1 | CP001037 - PAS/PAC sensor hybrid histidine kinase | 134.42 | 2.75E-33 | 40.40 | 40.40 | 880 | 1569 | 384 | 608 | 29.84% |
| Npun_F3797 | ACC82181.1 | CP001037 - multi-sensor signal transduction histidine kinase | 106.686 | 2.10E-24 | 32.30 | 32.30 | 1582 | 2244 | 384 | 608 | 29.84% |
| Npun_F5043 | ACC83382.1 | CP001037 - multi-sensor signal transduction histidine kinase | 150.984 | 1.80E-39 | 36.30 | 36.30 | 817 | 1521 | 384 | 608 | 29.84% |
| Npun_F2908 | ACC81440.1 | CP001037 - multi-sensor hybrid histidine kinase | 135.191 | 4.66E-33 | 34.50 | 34.50 | 1054 | 1719 | 384 | 608 | 29.84% |
| Npun_R4769 | ACC83120.1 | CP001037 - multi-component transcriptional regulator, winged helix family | 166.007 | 4.60E-43 | 41.30 | 41.30 | 2152 | 2814 | 384 | 608 | 29.84% |
| Npun_F1330 | ACC80042.1 | CP001037 - histidine kinase | 132.88 | 3.86E-34 | 37.70 | 37.70 | 496 | 1179 | 386 | 610 | 29.84% |
| Npun_R6203 | ACC84489.1 | CP001037 - integral membrane sensor signal transduction histidine kinase | 92.4337 | 1.68E-20 | 30.90 | 30.90 | 571 | 1224 | 386 | 608 | 29.58% |
| Npun_F5193 | ACC83525.1 | CP001037 - integral membrane sensor signal transduction histidine kinase | 96.6709 | 1.31E-21 | 29.80 | 29.80 | 694 | 1467 | 386 | 608 | 29.58% |
| Npun_F0303 | ACC79088.1 | CP001037 - integral membrane sensor signal transduction histidine kinase | 100.138 | 2.95E-23 | 29.30 | 29.30 | 412 | 1080 | 386 | 608 | 29.58% |
| Npun_R6227 | ACC84511.1 | CP001037 - integral membrane sensor signal transduction histidine kinase | 123.25 | 4.81E-31 | 38.20 | 38.20 | 433 | 1080 | 386 | 608 | 29.58% |
| Npun_F1277 | ACC79998.1 | CP001037 - PAS/PAC sensor signal transduction histidine kinase | 114.39 | 6.60E-27 | 35.00 | 35.00 | 1288 | 1935 | 384 | 605 | 29.44% |
| Npun_R3052 | ACC81547.1 | CP001037 - integral membrane sensor signal transduction histidine kinase | 135.961 | 1.40E-34 | 35.40 | 35.40 | 766 | 1410 | 387 | 608 | 29.44% |
| Npun_R3716 | ACC82106.1 | CP001037 - multi-sensor signal transduction histidine kinase | 112.079 | 3.83E-26 | 29.80 | 29.80 | 1189 | 1860 | 384 | 603 | 29.18% |
| Npun_R2485 | ACC81046.1 | CP001037 - histidine kinase | 118.627 | 1.40E-28 | 35.90 | 35.90 | 892 | 1539 | 386 | 605 | 29.18% |
| Npun_F3797 | ACC82181.1 | CP001037 - multi-sensor signal transduction histidine kinase | 97.8265 | 1.38E-21 | 31.00 | 31.00 | 94 | 744 | 152 | 370 | 29.05% |
| Npun_R5113 | ACC83447.1 | CP001037 - GAF sensor signal transduction histidine kinase | 82.0333 | 1.46E-16 | 30.60 | 30.60 | 1123 | 1713 | 163 | 371 | 27.72% |
| Npun_F1553 | ACC80242.1 | CP001037 - diguanylate cyclase with PAS/PAC and GAF sensors | 89.3521 | 6.62E-19 | 30.10 | 30.10 | 1573 | 2142 | 152 | 360 | 27.72% |
| Npun_R1060 | ACC79786.1 | CP001037 - diguanylate cyclase/phosphodiesterase with GAF sensor | 105.145 | 7.49E-24 | 34.50 | 34.50 | 1135 | 1689 | 158 | 360 | 26.92% |
| Npun_R6125 | ACC84413.1 | CP001037 - multi-sensor signal transduction histidine kinase | 86.2705 | 6.36E-18 | 29.10 | 29.10 | 463 | 993 | 159 | 354 | 25.99% |
| Npun_R1597 | ACC80282.1 | CP001037 - GAF sensor signal transduction histidine kinase | 82.0333 | 1.43E-16 | 31.50 | 31.50 | 85 | 621 | 165 | 359 | 25.86% |
| Npun_R3784 | ACC82169.1 | CP001037 - multi-sensor hybrid histidine kinase | 93.5893 | 3.83E-20 | 32.30 | 32.30 | 76 | 591 | 162 | 355 | 25.73% |
| Npun_R3572 | ACC81968.1 | CP001037 - multi-sensor hybrid histidine kinase | 90.1225 | 4.73E-19 | 32.80 | 32.80 | 421 | 981 | 167 | 353 | 24.80% |
