## Supplemental Table S4 for "Diel expression dynamics in filamentous cyanobacteria"

**Supplemental Table S4.** tBLASTn results for cyanobacterial CikA-like proteins.

| Locus Tag | Accession | Name | Bit-Score | E Value | % Pairwise Identity | % Identical Sites | Hit Start | Hit End | Query Start | Query End | % Query Coverage |
|---|---|---|---|---|---|---|---|---|---|---|---|
| Npun_F1000 | ACC79731.1 | CP001037 - GAF sensor signal transduction histidine kinase | 800.045 | 0.00E+00 | 70.50 | 70.30 | 1 | 2040 | 95 | 697 | 71.11 |
| Npun_F2363 | ACC80931.1 | CP001037 - multi-sensor hybrid histidine kinase | 159.458 | 1.66E-40 | 29.80 | 29.40 | 1411 | 3321 | 247 | 830 | 68.87 |
| Npun_R1685 | ACC80356.1 | CP001037 - multi-sensor hybrid histidine kinase | 222.631 | 8.97E-61 | 32.80 | 32.50 | 2083 | 3792 | 248 | 830 | 68.75 |
| Npun_R3691 | ACC82081.1 | CP001037 - multi-sensor hybrid histidine kinase | 230.72 | 2.03E-63 | 33.40 | 33.10 | 3556 | 5364 | 247 | 828 | 68.63 |
| Npun_F2854 | ACC81387.1 | CP001037 - GAF sensor signal transduction histidine kinase | 116.316 | 4.77E-27 | 26.50 | 26.40 | 1681 | 3144 | 236 | 692 | 53.89 |
| Npun_R5897 | ACC84193.1 | CP001037 - multi-sensor hybrid histidine kinase | 206.068 | 2.45E-55 | 31.40 | 31.30 | 3055 | 4458 | 379 | 829 | 53.18 |
| Npun_F6362 | ACC84634.1 | CP001037 - multi-sensor signal transduction histidine kinase | 146.362 | 1.10E-36 | 28.00 | 27.80 | 1276 | 2544 | 247 | 694 | 52.83 |
| Npun_R6149 | ACC84435.1 | CP001037 - multi-sensor signal transduction histidine kinase | 92.0485 | 1.39E-19 | 25.00 | 24.70 | 1210 | 2586 | 247 | 692 | 52.59 |
| Npun_R5149 | ACC83481.1 | CP001037 - GAF sensor signal transduction histidine kinase | 107.842 | 1.04E-24 | 26.90 | 26.60 | 712 | 1959 | 250 | 692 | 52.24 |
| Npun_R4776 | ACC83125.1 | CP001037 - multi-sensor signal transduction histidine kinase | 102.834 | 6.86E-23 | 25.80 | 25.60 | 1882 | 3285 | 251 | 692 | 52.12 |
| Npun_R2903 | ACC81436.1 | CP001037 - multi-sensor signal transduction histidine kinase | 160.614 | 6.82E-41 | 29.90 | 29.70 | 2089 | 3330 | 250 | 691 | 52.12 |
| Npun_R5113 | ACC83447.1 | CP001037 - GAF sensor signal transduction histidine kinase | 95.5153 | 1.09E-20 | 26.10 | 25.90 | 1684 | 3009 | 252 | 693 | 52.12 |
| Npun_R1597 | ACC80282.1 | CP001037 - GAF sensor signal transduction histidine kinase | 110.153 | 3.69E-25 | 28.00 | 27.80 | 1726 | 3108 | 252 | 692 | 52.00 |
| Npun_F5479 | ACC83786.1 | CP001037 - integral membrane sensor hybrid histidine kinase | 197.208 | 6.04E-53 | 38.30 | 38.10 | 1333 | 2496 | 446 | 820 | 44.22 |
| Npun_F5679 | ACC83981.1 | CP001037 - multi-sensor hybrid multi-kinase | 162.54 | 2.87E-41 | 33.20 | 33.10 | 5992 | 7107 | 468 | 830 | 42.81 |
| Npun_R1868 | ACC80525.1 | CP001037 - multi-sensor hybrid histidine kinase | 173.326 | 1.13E-45 | 33.20 | 33.00 | 1018 | 2133 | 468 | 830 | 42.81 |
| Npun_R6464 | ACC84732.1 | CP001037 - multi-sensor hybrid histidine kinase | 172.17 | 4.05E-45 | 34.60 | 34.40 | 1165 | 2271 | 468 | 830 | 42.81 |
| Npun_F5092 | ACC83426.1 | CP001037 - multi-sensor hybrid histidine kinase | 157.532 | 5.95E-40 | 30.50 | 30.30 | 2512 | 3750 | 468 | 830 | 42.81 |
| Npun_F2889 | ACC81422.1 | CP001037 - CBS sensor hybrid histidine kinase | 202.986 | 1.41E-54 | 32.80 | 32.40 | 1165 | 2373 | 468 | 830 | 42.81 |
| Npun_R4211 | ACC82586.1 | CP001037 - PAS/PAC sensor hybrid histidine kinase | 178.333 | 1.44E-46 | 34.50 | 34.30 | 2194 | 3309 | 468 | 829 | 42.69 |
| Npun_R2035 | ACC80662.1 | CP001037 - PAS/PAC sensor hybrid histidine kinase | 169.088 | 1.03E-43 | 34.30 | 34.10 | 2143 | 3237 | 468 | 829 | 42.69 |
| Npun_R1798 | ACC80460.1 | CP001037 - multi-sensor hybrid histidine kinase | 183.726 | 2.31E-48 | 36.50 | 36.30 | 1876 | 2994 | 468 | 829 | 42.69 |
| Npun_R6347 | ACC84619.1 | CP001037 - multi-sensor hybrid histidine kinase | 169.474 | 3.96E-44 | 34.20 | 34.00 | 1018 | 2142 | 468 | 829 | 42.69 |
| Npun_R0896 | ACC79633.1 | CP001037 - multi-sensor hybrid histidine kinase | 162.155 | 2.21E-41 | 30.50 | 30.30 | 1618 | 2811 | 468 | 829 | 42.69 |
| Npun_F2908 | ACC81440.1 | CP001037 - multi-sensor hybrid histidine kinase | 156.762 | 1.34E-39 | 31.60 | 31.50 | 3127 | 4239 | 468 | 829 | 42.69 |
| Npun_R3548 | ACC81949.1 | CP001037 - multi-sensor hybrid histidine kinase | 143.665 | 1.94E-35 | 31.30 | 31.10 | 3103 | 4392 | 468 | 829 | 42.69 |
| Npun_R2263 | ACC80852.1 | CP001037 - hybrid histidine kinase | 167.162 | 1.44E-43 | 33.20 | 32.70 | 649 | 1758 | 468 | 829 | 42.69 |
| Npun_DR038 | ACC85457.1 | CP001041 - putative PAS/PAC sensor protein | 179.489 | 6.34E-47 | 34.00 | 33.90 | 2782 | 3885 | 468 | 828 | 42.57 |
| Npun_R2268 | ACC80857.1 | CP001037 - PAS/PAC sensor hybrid histidine kinase | 190.274 | 2.01E-51 | 35.80 | 35.60 | 1033 | 2127 | 468 | 828 | 42.57 |
| Npun_F6350 | ACC84622.1 | CP001037 - PAS/PAC sensor hybrid histidine kinase | 173.326 | 3.32E-45 | 33.30 | 33.20 | 1525 | 2631 | 468 | 828 | 42.57 |
| Npun_R4744 | ACC83096.1 | CP001037 - PAS/PAC sensor hybrid histidine kinase | 170.244 | 6.10E-44 | 33.10 | 32.80 | 1948 | 3108 | 468 | 828 | 42.57 |
| Npun_F3541 | ACC81945.1 | CP001037 - PAS/PAC sensor hybrid histidine kinase | 167.162 | 9.06E-44 | 34.60 | 34.50 | 880 | 2025 | 468 | 828 | 42.57 |
| Npun_F2908 | ACC81440.1 | CP001037 - multi-sensor hybrid histidine kinase | 160.999 | 5.19E-41 | 29.80 | 29.60 | 1054 | 2277 | 468 | 828 | 42.57 |
| Npun_F1600 | ACC80285.1 | CP001037 - integral membrane sensor hybrid histidine kinase | 186.037 | 1.96E-50 | 37.30 | 37.20 | 490 | 1611 | 468 | 828 | 42.57 |
| Npun_F1211 | ACC79935.1 | CP001037 - integral membrane sensor hybrid histidine kinase | 181.415 | 8.90E-48 | 34.90 | 34.70 | 814 | 1941 | 468 | 828 | 42.57 |
| Npun_R1760 | ACC80422.1 | CP001037 - GAF sensor hybrid histidine kinase | 167.548 | 2.57E-44 | 34.80 | 34.70 | 661 | 1782 | 468 | 828 | 42.57 |
| Npun_R3591 | ACC81987.1 | CP001037 - multi-sensor hybrid histidine kinase | 196.438 | 1.64E-52 | 34.90 | 34.70 | 1987 | 3105 | 468 | 827 | 42.45 |
| Npun_R3572 | ACC81968.1 | CP001037 - multi-sensor hybrid histidine kinase | 182.956 | 7.12E-48 | 36.30 | 36.10 | 3388 | 4476 | 468 | 827 | 42.45 |
| Npun_F2346 | ACC80914.1 | CP001037 - multi-sensor hybrid histidine kinase | 162.54 | 1.75E-41 | 32.00 | 31.80 | 2611 | 3711 | 468 | 827 | 42.45 |
| Npun_R3083 | ACC81570.1 | CP001037 - multi-sensor hybrid histidine kinase | 154.451 | 1.46E-39 | 31.60 | 31.40 | 874 | 2082 | 468 | 827 | 42.45 |
| Npun_F4131 | ACC82509.1 | CP001037 - GAF sensor hybrid histidine kinase | 149.058 | 2.46E-38 | 31.40 | 31.20 | 559 | 1650 | 468 | 825 | 42.22 |
| Npun_R2262 | ACC80851.1 | CP001037 - PAS/PAC sensor hybrid histidine kinase | 160.229 | 1.50E-41 | 33.30 | 33.20 | 844 | 1953 | 468 | 824 | 42.10 |
| Npun_F5035 | ACC83378.1 | CP001037 - multi-sensor hybrid histidine kinase | 171.785 | 2.60E-44 | 32.70 | 32.50 | 4108 | 5313 | 468 | 824 | 42.10 |
| Npun_R3825 | ACC82209.1 | CP001037 - hybrid histidine kinase | 204.912 | 4.26E-57 | 37.80 | 37.60 | 697 | 1803 | 468 | 824 | 42.10 |
| Npun_R2375 | ACC80942.1 | CP001037 - multi-sensor hybrid histidine kinase | 205.297 | 3.41E-55 | 34.80 | 34.50 | 1582 | 2793 | 468 | 822 | 41.86 |
| Npun_R3784 | ACC82169.1 | CP001037 - multi-sensor hybrid histidine kinase | 95.1301 | 1.31E-20 | 27.80 | 27.60 | 1546 | 2628 | 475 | 829 | 41.86 |
| Npun_F2686 | ACC81226.1 | CP001037 - GAF sensor hybrid histidine kinase | 199.134 | 5.26E-53 | 34.60 | 34.40 | 4585 | 5748 | 468 | 820 | 41.63 |
| Npun_AR131 | ACC85002.1 | CP001038 - multi-sensor hybrid histidine kinase | 155.992 | 1.12E-39 | 34.60 | 34.40 | 1126 | 2202 | 468 | 816 | 41.16 |
| Npun_R4748 | ACC83100.1 | CP001037 - multi-sensor hybrid histidine kinase | 241.506 | 3.26E-69 | 42.60 | 42.40 | 1084 | 2175 | 468 | 816 | 41.16 |
| Npun_F1185 | ACC79910.1 | CP001037 - multi-sensor signal transduction histidine kinase | 137.117 | 2.45E-34 | 34.00 | 33.90 | 739 | 1677 | 364 | 692 | 38.80 |
| Npun_F5360 | ACC83673.1 | CP001037 - multi-sensor hybrid histidine kinase | 178.718 | 1.30E-46 | 39.20 | 39.00 | 2044 | 2991 | 468 | 771 | 35.85 |
| Npun_F6040 | ACC84330.1 | CP001037 - multi-sensor signal transduction histidine kinase | 172.17 | 6.97E-45 | 38.50 | 38.40 | 1792 | 2640 | 395 | 692 | 35.14 |
| Npun_R1012 | ACC79743.1 | CP001037 - integral membrane sensor signal transduction histidine kinase HepK | 161.384 | 2.36E-42 | 44.20 | 44.20 | 946 | 1707 | 446 | 692 | 29.13 |
| Npun_R1550 | ACC80239.1 | CP001037 - GAF sensor signal transduction histidine kinase | 135.961 | 8.06E-34 | 36.00 | 36.00 | 994 | 1716 | 468 | 696 | 27.00 |
| Npun_F0303 | ACC79088.1 | CP001037 - integral membrane sensor signal transduction histidine kinase | 120.553 | 4.31E-30 | 32.60 | 32.60 | 406 | 1089 | 468 | 695 | 26.89 |
| Npun_R5764 | ACC84062.1 | CP001037 - histidine kinase with KaiB domain, SasA | 93.9745 | 5.35E-21 | 30.10 | 30.10 | 505 | 1200 | 468 | 695 | 26.89 |
| Npun_BF140 | ACC85272.1 | CP001039 - integral membrane sensor signal transduction histidine kinase | 103.219 | 7.15E-24 | 31.40 | 31.30 | 574 | 1239 | 470 | 696 | 26.77 |
| Npun_R1449 | ACC80152.1 | CP001037 - response regulator receiver sensor signal transduction histidine kinase | 184.496 | 2.72E-51 | 44.00 | 44.00 | 520 | 1194 | 468 | 694 | 26.77 |
| Npun_R1448 | ACC80151.1 | CP001037 - response regulator receiver sensor signal transduction histidine kinase | 119.398 | 1.21E-29 | 30.00 | 30.00 | 424 | 1119 | 468 | 694 | 26.77 |
| Npun_R0454 | ACC79233.1 | CP001037 - GAF sensor signal transduction histidine kinase | 98.9821 | 3.30E-22 | 30.50 | 30.50 | 688 | 1518 | 468 | 694 | 26.77 |
| Npun_R3054 | ACC81549.1 | CP001037 - Chase sensor signal transduction histidine kinase | 118.627 | 6.27E-29 | 34.40 | 34.40 | 619 | 1284 | 468 | 694 | 26.77 |
| Npun_F0957 | ACC79688.1 | CP001037 - response regulator receiver sensor signal transduction histidine kinase | 119.783 | 8.57E-30 | 32.00 | 32.00 | 424 | 1077 | 468 | 692 | 26.53 |
| Npun_F0022 | ACC78825.1 | CP001037 - response regulator receiver sensor signal transduction histidine kinase | 116.316 | 1.89E-28 | 31.90 | 31.90 | 487 | 1149 | 468 | 692 | 26.53 |
| Npun_F0839 | ACC79577.1 | CP001037 - response regulator receiver sensor signal transduction histidine kinase | 113.62 | 1.36E-27 | 31.00 | 31.00 | 424 | 1128 | 468 | 692 | 26.53 |
| Npun_F4953 | ACC83298.1 | CP001037 - PAS/PAC sensor signal transduction histidine kinase | 129.028 | 2.86E-32 | 34.70 | 34.70 | 667 | 1326 | 468 | 692 | 26.53 |
| Npun_F1277 | ACC79998.1 | CP001037 - PAS/PAC sensor signal transduction histidine kinase | 120.168 | 1.34E-28 | 35.40 | 35.30 | 1288 | 1944 | 468 | 692 | 26.53 |
| Npun_R3198 | ACC81651.1 | CP001037 - PAS/PAC sensor signal transduction histidine kinase | 113.62 | 1.29E-27 | 34.50 | 34.50 | 457 | 1116 | 468 | 692 | 26.53 |
| Npun_F5043 | ACC83382.1 | CP001037 - multi-sensor signal transduction histidine kinase | 149.443 | 8.80E-39 | 36.40 | 36.40 | 817 | 1521 | 468 | 692 | 26.53 |
| Npun_F6002 | ACC84292.1 | CP001037 - multi-sensor signal transduction histidine kinase | 122.479 | 7.89E-30 | 33.70 | 33.60 | 817 | 1482 | 468 | 692 | 26.53 |
| Npun_F3797 | ACC82181.1 | CP001037 - multi-sensor signal transduction histidine kinase | 118.627 | 5.01E-28 | 35.00 | 34.90 | 1582 | 2244 | 468 | 692 | 26.53 |
| Npun_R6125 | ACC84413.1 | CP001037 - multi-sensor signal transduction histidine kinase | 85.1149 | 1.50E-17 | 29.30 | 29.30 | 1573 | 2247 | 468 | 692 | 26.53 |
| Npun_F2346 | ACC80914.1 | CP001037 - multi-sensor hybrid histidine kinase | 137.502 | 1.22E-33 | 39.40 | 39.40 | 1297 | 1962 | 468 | 692 | 26.53 |
| Npun_R4769 | ACC83120.1 | CP001037 - multi-component transcriptional regulator, winged helix family | 158.688 | 2.31E-40 | 41.70 | 41.70 | 2152 | 2814 | 468 | 692 | 26.53 |
| Npun_R6227 | ACC84511.1 | CP001037 - integral membrane sensor signal transduction histidine kinase | 132.109 | 5.44E-34 | 39.70 | 39.60 | 427 | 1080 | 468 | 692 | 26.53 |
| Npun_R1236 | ACC79960.1 | CP001037 - integral membrane sensor signal transduction histidine kinase | 135.961 | 1.61E-33 | 37.70 | 37.70 | 1153 | 1830 | 468 | 692 | 26.53 |
| Npun_F1330 | ACC80042.1 | CP001037 - histidine kinase | 133.265 | 3.99E-34 | 36.20 | 36.20 | 496 | 1179 | 470 | 694 | 26.53 |
| Npun_R1759 | ACC80421.1 | CP001037 - GAF sensor signal transduction histidine kinase | 133.65 | 8.52E-33 | 40.70 | 40.60 | 1393 | 2058 | 468 | 692 | 26.53 |
| Npun_F3565 | ACC81961.1 | CP001037 - multi-sensor signal transduction multi-kinase | 92.4337 | 1.47E-19 | 29.00 | 28.90 | 5500 | 6150 | 471 | 694 | 26.42 |
| Npun_R2485 | ACC81046.1 | CP001037 - histidine kinase | 101.679 | 5.37E-23 | 33.10 | 33.00 | 886 | 1545 | 468 | 691 | 26.42 |
| Npun_F0020 | ACC78823.1 | CP001037 - multi-sensor signal transduction histidine kinase | 87.4261 | 3.34E-18 | 31.20 | 31.10 | 1591 | 2235 | 470 | 692 | 26.30 |
| Npun_F1439 | ACC80146.1 | CP001037 - integral membrane sensor signal transduction histidine kinase | 133.265 | 1.38E-33 | 35.90 | 35.90 | 763 | 1413 | 470 | 692 | 26.30 |
| Npun_F5193 | ACC83525.1 | CP001037 - integral membrane sensor signal transduction histidine kinase | 111.694 | 2.48E-26 | 32.50 | 32.40 | 694 | 1467 | 470 | 692 | 26.30 |
| Npun_R6203 | ACC84489.1 | CP001037 - integral membrane sensor signal transduction histidine kinase | 100.523 | 4.58E-23 | 30.50 | 30.50 | 571 | 1224 | 470 | 692 | 26.30 |
| Npun_R3052 | ACC81547.1 | CP001037 - integral membrane sensor signal transduction histidine kinase | 133.265 | 1.33E-33 | 36.10 | 36.00 | 766 | 1410 | 471 | 692 | 26.18 |
| Npun_R3716 | ACC82106.1 | CP001037 - multi-sensor signal transduction histidine kinase | 121.324 | 4.61E-29 | 32.80 | 32.80 | 1189 | 1860 | 468 | 687 | 25.94 |
| Npun_F3675 | ACC82065.1 | CP001037 - multi-sensor signal transduction histidine kinase | 93.2041 | 2.55E-20 | 29.30 | 29.20 | 850 | 1479 | 471 | 687 | 25.59 |
| Npun_R6125 | ACC84413.1 | CP001037 - multi-sensor signal transduction histidine kinase | 89.7373 | 5.26E-19 | 32.40 | 32.10 | 433 | 939 | 238 | 420 | 21.58 |
| Npun_F1553 | ACC80242.1 | CP001037 - diguanylate cyclase with PAS/PAC and GAF sensors | 97.8265 | 2.23E-21 | 36.90 | 36.80 | 1573 | 2070 | 241 | 420 | 21.23 |
| Npun_R3784 | ACC82169.1 | CP001037 - multi-sensor hybrid histidine kinase | 111.694 | 1.11E-25 | 39.90 | 39.50 | 43 | 534 | 244 | 420 | 20.87 |
| Npun_F3797 | ACC82181.1 | CP001037 - multi-sensor signal transduction histidine kinase | 89.3521 | 8.02E-19 | 33.40 | 33.30 | 106 | 642 | 245 | 420 | 20.75 |
| Npun_R1597 | ACC80282.1 | CP001037 - GAF sensor signal transduction histidine kinase | 93.5893 | 5.20E-20 | 33.60 | 33.50 | 67 | 558 | 247 | 422 | 20.75 |
| Npun_R5113 | ACC83447.1 | CP001037 - GAF sensor signal transduction histidine kinase | 97.8265 | 2.45E-21 | 35.00 | 34.90 | 55 | 537 | 251 | 422 | 20.28 |
| Npun_F1883 | ACC80538.1 | CP001037 - putative GAF sensor protein | 95.5153 | 1.07E-20 | 37.90 | 37.50 | 1504 | 1986 | 251 | 420 | 20.05 |
| Npun_F1883 | ACC80538.1 | CP001037 - putative GAF sensor protein | 89.7373 | 6.90E-19 | 35.80 | 35.40 | 2020 | 2502 | 251 | 420 | 20.05 |
| Npun_R4776 | ACC83125.1 | CP001037 - multi-sensor signal transduction histidine kinase | 90.1225 | 6.14E-19 | 32.90 | 32.60 | 1327 | 1815 | 251 | 420 | 20.05 |
| Npun_R3572 | ACC81968.1 | CP001037 - multi-sensor hybrid histidine kinase | 90.8929 | 3.98E-19 | 31.30 | 31.20 | 406 | 930 | 251 | 420 | 20.05 |
| Npun_R1060 | ACC79786.1 | CP001037 - diguanylate cyclase/phosphodiesterase with GAF sensor | 127.102 | 1.25E-30 | 43.10 | 42.70 | 1147 | 1617 | 251 | 420 | 20.05 |
| Npun_R5113 | ACC83447.1 | CP001037 - GAF sensor signal transduction histidine kinase | 85.5001 | 1.61E-17 | 33.10 | 32.90 | 1129 | 1599 | 254 | 421 | 19.81 |
| Npun_AF142 | ACC85008.1 | CP001038 - GAF sensor signal transduction histidine kinase | 94.7449 | 1.98E-20 | 35.30 | 35.20 | 2161 | 2625 | 256 | 420 | 19.46 |
| Npun_R5313 | ACC83634.1 | CP001037 - multi-sensor signal transduction histidine kinase | 88.1965 | 1.82E-18 | 33.20 | 33.10 | 1315 | 1755 | 259 | 421 | 19.22 |
