## Supplemental Table S5 for "Diel expression dynamics in filamentous cyanobacteria"

Supplemental Table S5. Putative CikA homologue expression correlation.

| Gene | CikA-like_Gene | CorrelationValue |
|---|---|---|
| Npun_R1733 (LMW PBP) | Npun_F2854 | 0.982449443 |
| Npun_R1733 (LMW PBP) | Npun_R5113 | 0.971446745 |
| Npun_R1952 (UppS) | Npun_R1685 | 0.968856927 |
| Npun_R1733 (LMW PBP) | Npun_R1685 | 0.968649658 |
| Npun_R1733 (LMW PBP) | Npun_F2363 | 0.954675064 |
| Npun_R1733 (LMW PBP) | Npun_R1597 | 0.949939303 |
| Npun_F5138 (FtsE) | Npun_R5113 | 0.94517937 |
| Npun_F5138 (FtsE) | Npun_F2363 | 0.938532519 |
| Npun_R1839 (MreD) | Npun_R5113 | 0.927348683 |
| Npun_R1952 (UppS) | Npun_F2363 | 0.921459773 |
| Npun_R1952 (UppS) | Npun_R5113 | 0.916957407 |
| Npun_R0056 (DUF152) | Npun_R6149 | 0.916119909 |
| Npun_R4092 (FtsK) | Npun_F6362 | 0.914128122 |
| Npun_F5138 (FtsE) | Npun_R1597 | 0.911793486 |
| Npun_R1841 (MreB) | Npun_R5113 | 0.911662811 |
| Npun_R1839 (MreD) | Npun_R1597 | 0.908563261 |
| Npun_R1952 (UppS) | Npun_F2854 | 0.905274232 |
| Npun_F3659 (RpaA) | Npun_R3691 | 0.899298892 |
| Npun_F5138 (FtsE) | Npun_F2854 | 0.892876802 |
| Npun_R1841 (MreB) | Npun_F2854 | 0.886233362 |
| Npun_F4881 (FtsH) | Npun_F6362 | 0.885568809 |
| Npun_R1839 (MreD) | Npun_F2854 | 0.885381621 |
| Npun_F5138 (FtsE) | Npun_R1685 | 0.881389818 |
| Npun_F3647 (MinC) | Npun_R3691 | 0.880990067 |
| Npun_R1302 (NagB) | Npun_R1597 | 0.87842755 |
| Npun_R4933 (Cdv1) | Npun_R5149 | 0.875761242 |
| Npun_R4507 (BacA) | Npun_R5149 | 0.875678492 |
| Npun_F5214 (GlmS) | Npun_R6149 | 0.875275584 |
| Npun_F2411 (MurG) | Npun_R5149 | 0.875162279 |
| Npun_R1302 (NagB) | Npun_F2363 | 0.873216278 |
| Npun_R1840 (MreC) | Npun_R1597 | 0.870303677 |
| Npun_R1839 (MreD) | Npun_F2363 | 0.868200199 |
| Npun_R1840 (MreC) | Npun_R5113 | 0.864124214 |
| Npun_R3910 (BolA) | Npun_F6362 | 0.855022767 |
| Npun_R1302 (NagB) | Npun_F2854 | 0.854136988 |
| Npun_R1841 (MreB) | Npun_R1597 | 0.84607166 |
| Npun_F3659 (RpaA) | Npun_F6362 | 0.840567752 |
| Npun_R1302 (NagB) | Npun_R4776 | 0.837799452 |
| Npun_R1952 (UppS) | Npun_R1597 | 0.837440445 |

| | | |
|---|---|---|
| Npun_R6629 (SepI) | Npun_R3691 | 0.836204771 |
| Npun_R1840 (MreC) | Npun_F2854 | 0.832754335 |
| Npun_R4455 (VanY) | Npun_F6362 | 0.832644182 |
| Npun_F5597 (Ddl) | Npun_F6362 | 0.831635338 |
| Npun_R4092 (FtsK) | Npun_R3691 | 0.829762013 |
| Npun_F0907 (GlmU) | Npun_R5149 | 0.82564693 |
| Npun_R1302 (NagB) | Npun_R1685 | 0.817145904 |
| Npun_F3647 (MinC) | Npun_F6362 | 0.815084713 |
| Npun_F5597 (Ddl) | Npun_R3691 | 0.812478151 |
| Npun_F3647 (MinC) | Npun_R6149 | 0.808768838 |
| Npun_R1839 (MreD) | Npun_R1685 | 0.806995872 |
| Npun_R1302 (NagB) | Npun_R5113 | 0.805592872 |
| Npun_R4455 (VanY) | Npun_R3691 | 0.796933605 |
| Npun_R4933 (Cdv1) | Npun_R2903 | 0.795042785 |
| Npun_R0056 (DUF152) | Npun_R3691 | 0.793707855 |
| Npun_F3659 (RpaA) | Npun_R6149 | 0.786853935 |
| Npun_F5950 (YlmG) | Npun_R2903 | 0.786809327 |
| Npun_F5950 (YlmG) | Npun_F1000 | 0.786164113 |
| Npun_R6354 (YlmH) | Npun_R3691 | 0.781211921 |
| Npun_R1841 (MreB) | Npun_R1685 | 0.780374542 |
| Npun_F4453 (Class B PBP) | Npun_R5149 | 0.773071614 |
| Npun_R0056 (DUF152) | Npun_F6362 | 0.762959683 |
| Npun_F4452 (Class B PBP) | Npun_F2854 | 0.760014664 |
