## Supplemental Table S6 for "Diel expression dynamics in filamentous cyanobacteria"

Supplemental Table S6. Loci associated with motility and/or hormogonia development.

| Locus Tag | Gene Name | Function | Mutant Phenotype |
|---|---|---|---|
| Npun_F0676 | pilA | major pilin | non-motile |
| Npun_R0116 | pilC | innermembrane platform | non-motile |
| Npun_R0117 | pilT1 | pilus retraction ATPase | non-motile, hyperpiliated |
| Npun_R0118 | pilB | pilus extension ATPase | non-motile, loss of pilus extension |
| Npun_F2507 | pilT2 | pilus retraction ATPase | reduced motility |
| Npun_F5005 | pilM | pilus alignment complex protein | non-motile |
| Npun_F5006 | pilN | pilus alignment complex protein | |
| Npun_F5007 | pilO | pilus alignment complex protein | |
| Npun_F5008 | pilQ | outermembrane secretin | non-motile |
| Npun_F5230 | hfq | Hfq homolog, essential for PilB activity | non-motile, loss of pilus extension |
| Npun_F4125 | ebsA | polysaccharide biosynthesis | non-motile |
| Npun_F0677 | ogtA | O-linked-B-N-acetylglucosamine transferase | non-motile, fails to accumulate PilA |
| Npun_F0066 | hpsA | conserved hypothetical membrane protein | non-motile, reduced HPS |
| Npun_F0067 | hpsB | minor pilin | non-motile, reduced HPS |
| Npun_F0068 | hpsC | minor pilin | non-motile, reduced HPS |
| Npun_F0069 | hpsD | minor pilin | non-motile, reduced HPS |
| Npun_F0070 | hpsE | glycosyl transferase | non-motile, HPS- |
| Npun_F0071 | hpsF | glycosyl transferase | non-motile, HPS- |
| Npun_F0072 | hpsG | glycosyl transferase | non-motile, HPS- |
| Npun_F0073 | hpsH | minor pilin | |
| Npun_F0075 | hpsI | glycosyl transferase | |
| Npun_F0077 | hpsJ | conserved hypothetical membrane protein | reduced motility, altered HPS composition |
| Npun_F0078 | hpsK | glycosyl transferase | |
| Npun_R0640 | hpsL | O-antigen ligase-like membrane protein | non-motile, HPS- |
| Npun_R0639 | hpsM | glycosyl transferase | reduced motiltiy |
| Npun_R0638 | hpsN | glycosyl transferase | non-motile, HPS- |
| Npun_R0637 | hpsO | glycosyl transferase | non-motile, HPS- |
| Npun_R0636 | hpsP | glycosyl transferase | non-motile, HPS- |
| Npun_F1388 | hpsQ | glycosyl transferase | non-motile, HPS- |
| Npun_R1506 | hpsR | glycosyl transferase | non-motile, HPS- |
| Npun_R5614 | hpsS | glycosyl transferase | non-motile, HPS- |
| Npun_R5613 | hpsT | glycosyl transferase | |
| Npun_R6512 | hpsU | WcaF | |
| Npun_R0453 | wzy | polysaccharide polymerase | |
| Npun_F0458 | wza | polysaccharide export outer membrane protein | |
| Npun_F0459 | wzc | polysaccharide co-polymerase | |
| Npun_F5960 | hmpA | PatA-type response regulator | motile |
| Npun_F5961 | hmpB | CheY-type response regulator | non-motile, reduced HPS |
| Npun_F5962 | hmpC | CheW | non-motile, reduced HPS |
| Npun_F5963 | hmpD | MCP | non-motile, reduced HPS |
| Npun_F5964 | hmpE | CheA | non-motile, reduced HPS |
| Npun_R5959 | hmpF | coiled-coil protein | non-motile, loss of pilus extension |
| Npun_F2161 | ptxA | CheY, PatA family | |
| Npun_F2162 | ptxB | CheY | |
| Npun_F2163 | ptxC | CheW | |
| Npun_F2164 | ptxD | MCP | motile, loss of phototaxis |
| Npun_F2165 | ptxE | CheA | motile, loss of phototaxis |

| | | | |
|---|---|---|---|
| Npun_F2166 | ptxF | CheW | |
| Npun_F2167 | ptxG | MCP fragment | |
| Npun_F2168 | ptxH | CheA fragment | motile, reduced phototaxis |
| Npun_R5135 | hmpU | RsbU-type phosphatase | reduced motility/hormognoium development |
| Npun_R5134 | hmpW | RsbW-type kinase/anti-sigma factor | hyper motile/enhanced hormogonium developm[illegible] |
| Npun_F5169 | hmpV | RsbV-type anti-sigma antagonist | reduced motility/hormognoium development |
| Npun_F1682 | hcyA | RsbV-type anti-sigma antagonist | reduced motility |
| Npun_R1683 | hcyB | RsbU-type phosphatase | |
| Npun_F1684 | hcyC | RsbW-type kinase/anti-sigma factor | |
| Npun_R1685 | hcyD | Histidine kinase, GAF sensor | |
| Npun_R3825 | hrmX (formerly hrmK) | hybrid histidine kinase/response regulator | Hormogonium- |
| Npun_R1337 | sigJ | alternative sigma factor | Hormogonium- |
| Npun_F0996 | sigC | alternative sigma factor | Hormogonium- |
| Npun_F4811 | sigF | alternative sigma factor | non-motile, loss of pilA expression |
| Npun_F0122 | dnaK1 | DnaK/Hsp70 type chaperone | reduced motility/HPS |
| Npun_F1160 | dnaJ3 | DnaJ co-chaperone | reduced motility/HPS |
