## Supplemental Table S7 for "Diel expression dynamics in filamentous cyanobacteria"

Supplemental Table S7. Expression correlation matrix for DGR-associated genes.

| | Npun_F4884 | Npun_F4885 | Npun_F4886 | Npun_F4887 | Npun_F4888 | **Npun_F4889** | **Npun_F4890** | **Npun_F4891** | **Npun_F4892** | **DGR-TR** | Npun_F4893 | Npun_F4894 | Npun_F4895 | Npun_F4896 | Npun_F4897 | Annotation |
|---|---|---|---|---|---|---|---|---|---|---|---|---|---|---|---|---|
| Npun_F4884 | 1.0000 | 0.1827 | 0.3200 | 0.5340 | 0.3097 | -0.2250 | 0.0000 | 0.7209 | 0.0637 | -0.4366 | 0.3529 | 0.2061 | -0.3143 | 0.0880 | -0.2436 | Hypothetical protein |
| Npun_F4885 | 0.1827 | 1.0000 | 0.8855 | 0.8370 | 0.6940 | -0.1011 | 0.7616 | 0.2538 | 0.1663 | 0.2213 | 0.3876 | -0.0107 | 0.0355 | 0.3152 | -0.1244 | CHAT domain protein |
| Npun_F4886 | 0.3200 | 0.8855 | 1.0000 | 0.8116 | 0.9064 | 0.2296 | 0.7382 | 0.3127 | 0.4173 | -0.0526 | 0.5285 | -0.2514 | 0.0305 | 0.6137 | 0.0696 | Hypothetical protein |
| Npun_F4887 | 0.5340 | 0.8370 | 0.8116 | 1.0000 | 0.7205 | -0.0777 | 0.5562 | 0.3386 | 0.1605 | 0.1880 | 0.4294 | -0.0774 | 0.0242 | 0.4239 | -0.1876 | MoxR ATPase |
| Npun_F4888 | 0.3097 | 0.6940 | 0.9064 | 0.7205 | 1.0000 | 0.4065 | 0.7399 | 0.3002 | 0.6518 | -0.0192 | 0.6820 | -0.3750 | 0.0369 | 0.7537 | 0.3606 | VWA protein |
| **Npun_F4889** | -0.2250 | -0.1011 | 0.2296 | -0.0777 | 0.4065 | 1.0000 | 0.1874 | -0.4863 | 0.2871 | -0.2454 | 0.3552 | -0.5761 | 0.2788 | 0.7734 | 0.3725 | **DGR-VP1** |
| **Npun_F4890** | 0.0000 | 0.7616 | 0.7382 | 0.5562 | 0.7399 | 0.1874 | 1.0000 | 0.0713 | 0.3338 | 0.1903 | 0.5653 | -0.1639 | 0.0504 | 0.4925 | 0.2729 | **DGR-VP2** |
| **Npun_F4891** | 0.7209 | 0.2538 | 0.3127 | 0.3386 | 0.3002 | -0.4863 | 0.0713 | 1.0000 | 0.3225 | -0.2638 | 0.3243 | 0.4269 | -0.4950 | -0.2210 | 0.0359 | **DGR-Avd** |
| **Npun_F4892** | 0.0637 | 0.1663 | 0.4173 | 0.1605 | 0.6518 | 0.2871 | 0.3338 | 0.3225 | 1.0000 | 0.1369 | 0.4499 | -0.5021 | -0.2648 | 0.3966 | 0.7839 | **DGR-RT** |
| **DGR-TR** | -0.4366 | 0.2213 | -0.0526 | 0.1880 | -0.0192 | -0.2454 | 0.1903 | -0.2638 | 0.1369 | 1.0000 | -0.0581 | -0.1546 | -0.1065 | -0.1759 | 0.2404 | **DGR-TR** |
| Npun_F4893 | 0.3529 | 0.3876 | 0.5285 | 0.4294 | 0.6820 | 0.3552 | 0.5653 | 0.3243 | 0.4499 | -0.0581 | 1.0000 | -0.1006 | -0.4077 | 0.3684 | 0.4785 | HTH Regulator |
| Npun_F4894 | 0.2061 | -0.0107 | -0.2514 | -0.0774 | -0.3750 | -0.5761 | -0.1639 | 0.4269 | -0.5021 | -0.1546 | -0.1006 | 1.0000 | -0.0232 | -0.6033 | -0.4418 | Hypothetical protein |
| Npun_F4895 | -0.3143 | 0.0355 | 0.0305 | 0.0242 | 0.0369 | 0.2788 | 0.0504 | -0.4950 | -0.2648 | -0.1065 | -0.4077 | -0.0232 | 1.0000 | 0.4298 | -0.3552 | Hypothetical protein |
| Npun_F4896 | 0.0880 | 0.3152 | 0.6137 | 0.4239 | 0.7537 | 0.7734 | 0.4925 | -0.2210 | 0.3966 | -0.1759 | 0.3684 | -0.6033 | 0.4298 | 1.0000 | 0.1685 | Helicase |
| Npun_F4897 | -0.2436 | -0.1244 | 0.0696 | -0.1876 | 0.3606 | 0.3725 | 0.2729 | 0.0359 | 0.7839 | 0.2404 | 0.4785 | -0.4418 | -0.3552 | 0.1685 | 1.0000 | Restriction endonuclease |
| Annotation | Hypothetical protein | CHAT domain protein | Hypothetical protein | MoxR ATPase | VWA protein | **DGR-VP1** | **DGR-VP2** | **DGR-Avd** | **DGR-RT** | **DGR-TR** | HTH Regulator | Hypothetical protein | Hypothetical protein | Helicase | Restriction endonuclease | |
