## Supplemental Figure S1 for "Diel expression dynamics in filamentous cyanobacteria"

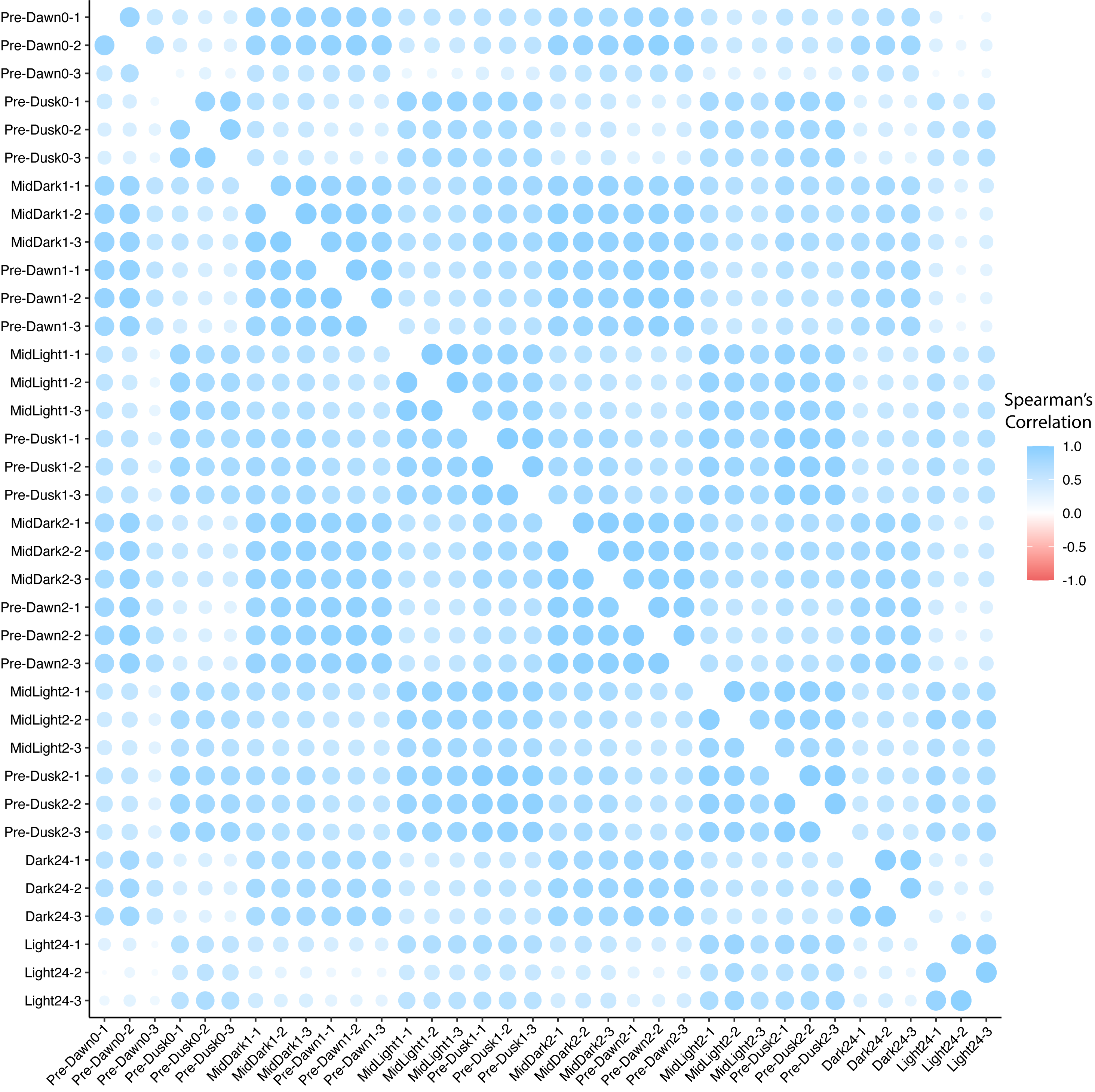


Supplemental Figure S1: Spearman correlation analysis of normalized expression counts.

Spearman correlation plot of normalized expression counts for all samples across the time-course experiment. Each circle represents the Spearman correlation coefficient between pairs of samples, with values ranging from -1.0 (perfect negative correlation) to 1.0 (perfect positive correlation). The size of each circle is proportional to the amount of transcriptional data. The heatmap is color-coded, with blue indicating positive correlations and red indicating negative correlations. Timepoints include Pre-Dawn, MidDark, Pre-Dusk, MidLight, Dark24, and Light24, with triplicate samples for each condition.
