## Supplemental Figure S2 for "Diel expression dynamics in filamentous cyanobacteria"

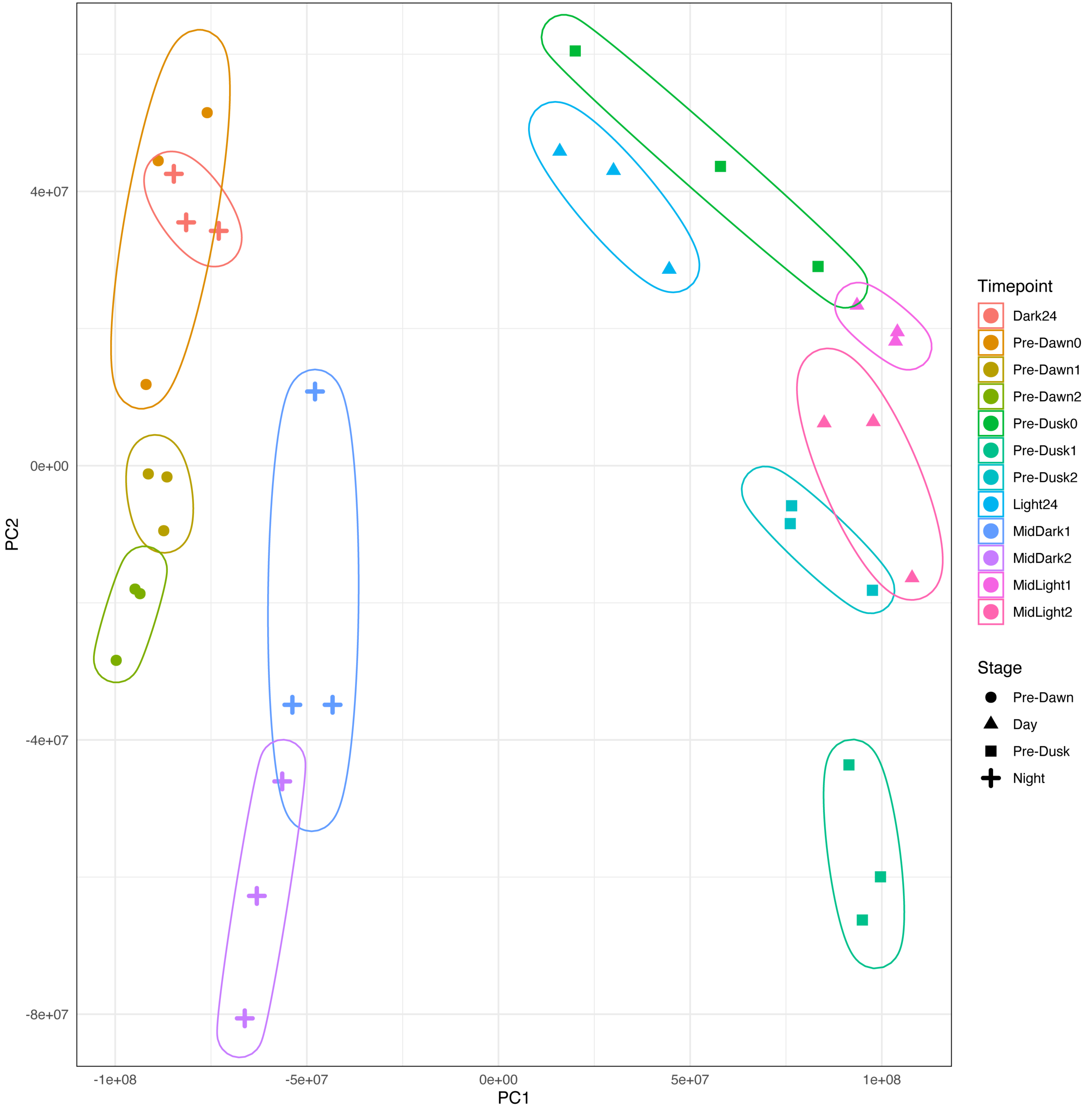


**Supplemental Figure S2.** Principal component analysis of normalized expression values.

Principal Component Analysis (PCA) of the normalized expression values for all 36 samples across the time-course experiment. Expression distribution by sample is shown for the first two principal components (PC1 and PC2). Each point represents a distinct sample, and timepoints are shown as separate colors. Shapes indicate the stages: circles for Pre-Dawn, squares for Pre-Dusk, triangles for Day, and plus-signs for Night.
