## Supplemental Figure S3 for "Diel expression dynamics in filamentous cyanobacteria"

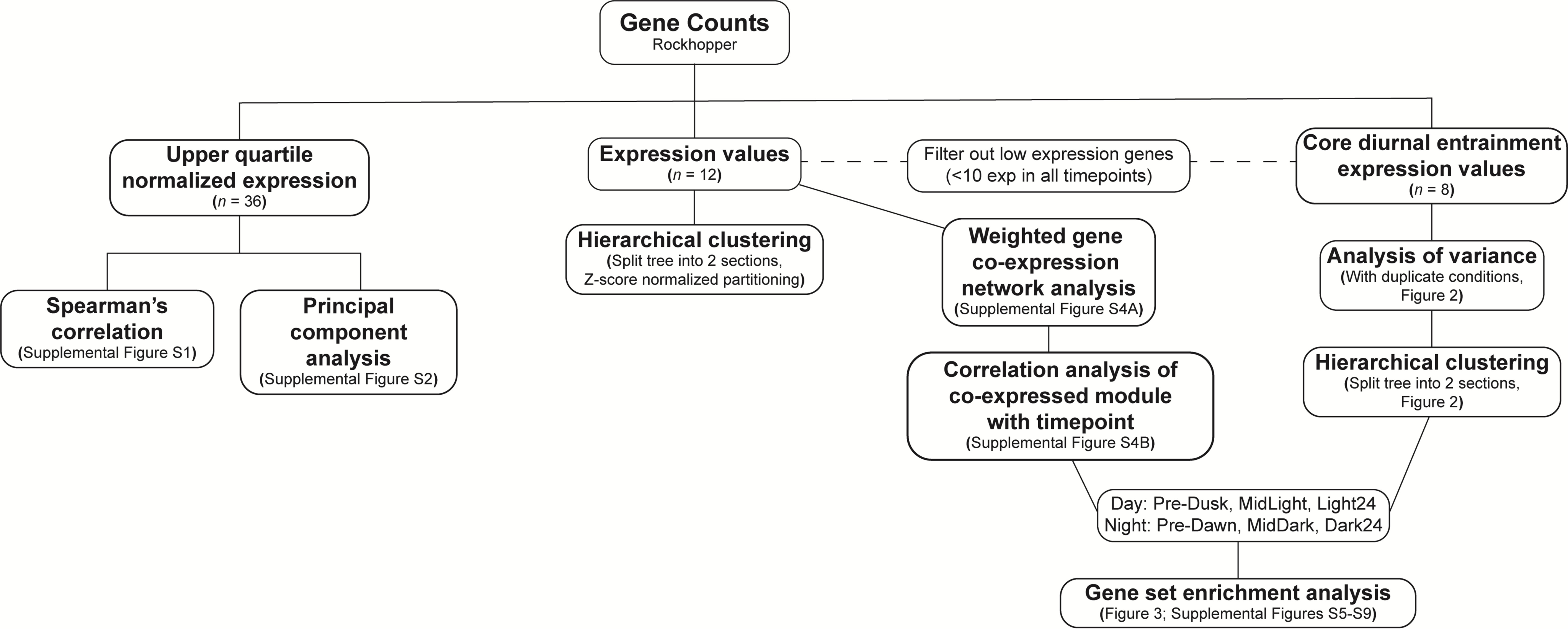


**Supplemental Figure S3.** Transcriptional data processing workflow.
Flowchart outlining the transcriptional data processing steps. Upper quartile normalized expression for all 36 samples was used for Spearman's correlation and principal component analysis. Low expression genes (less than 10 expression in all timepoints) were filtered out before subsequent analyses.
