## Supplemental Figure S4 for "Diel expression dynamics in filamentous cyanobacteria"

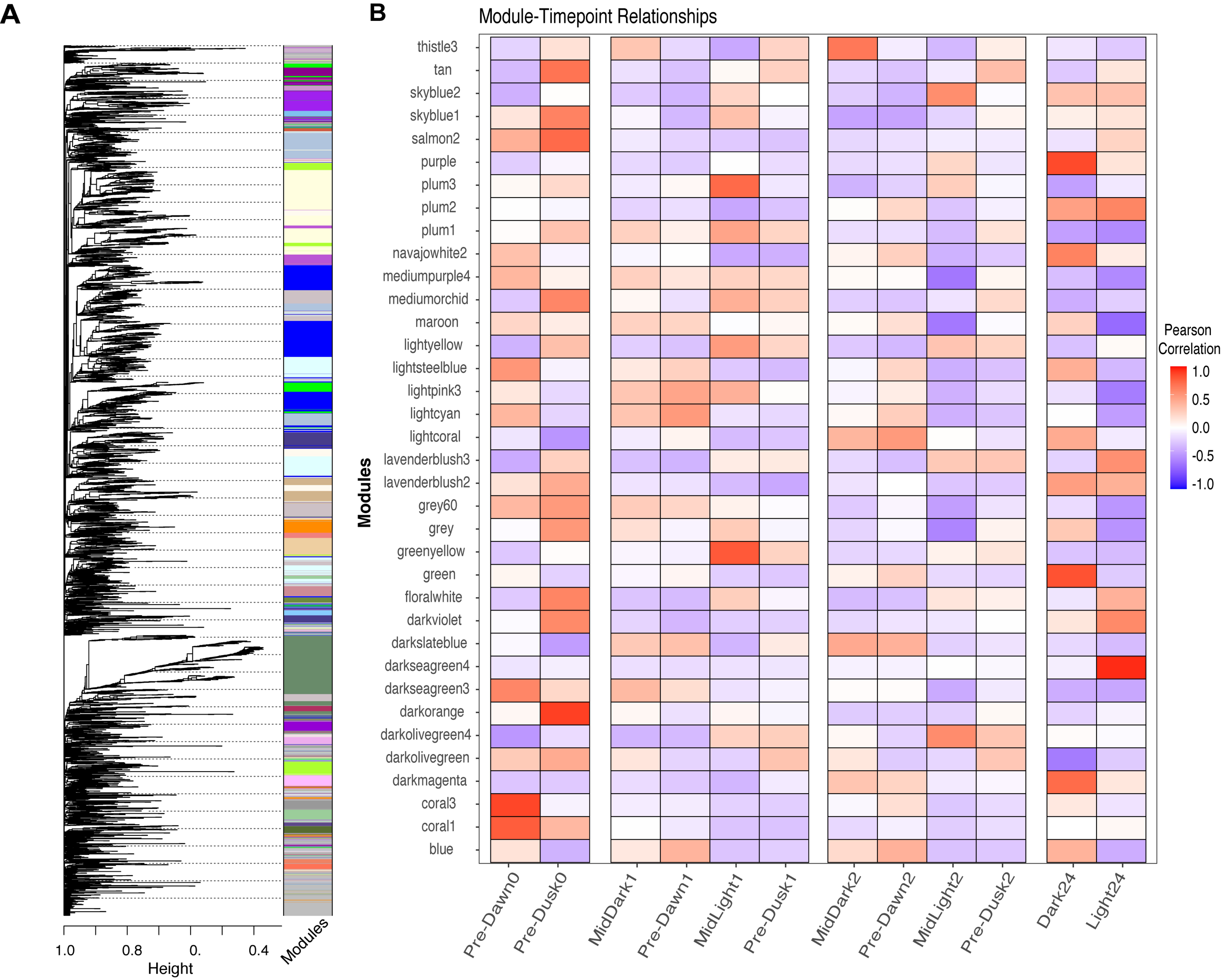


**Supplemental Figure S4.** Weighted gene co-expression network analysis.
(A) Clustering of the transcriptome into modules of co-expressed genes. Each branch represents a module containing genes with similar expression patterns across the 12 timepoints. (B) Pearson correlation matrix showing the relationships between the 12 timepoints and the modules of co-expressed genes. The correlation coefficients are color-coded, with blue indicating negative correlations and red indicating positive correlations.
