## Supplemental Figure S5 for "Diel expression dynamics in filamentous cyanobacteria"

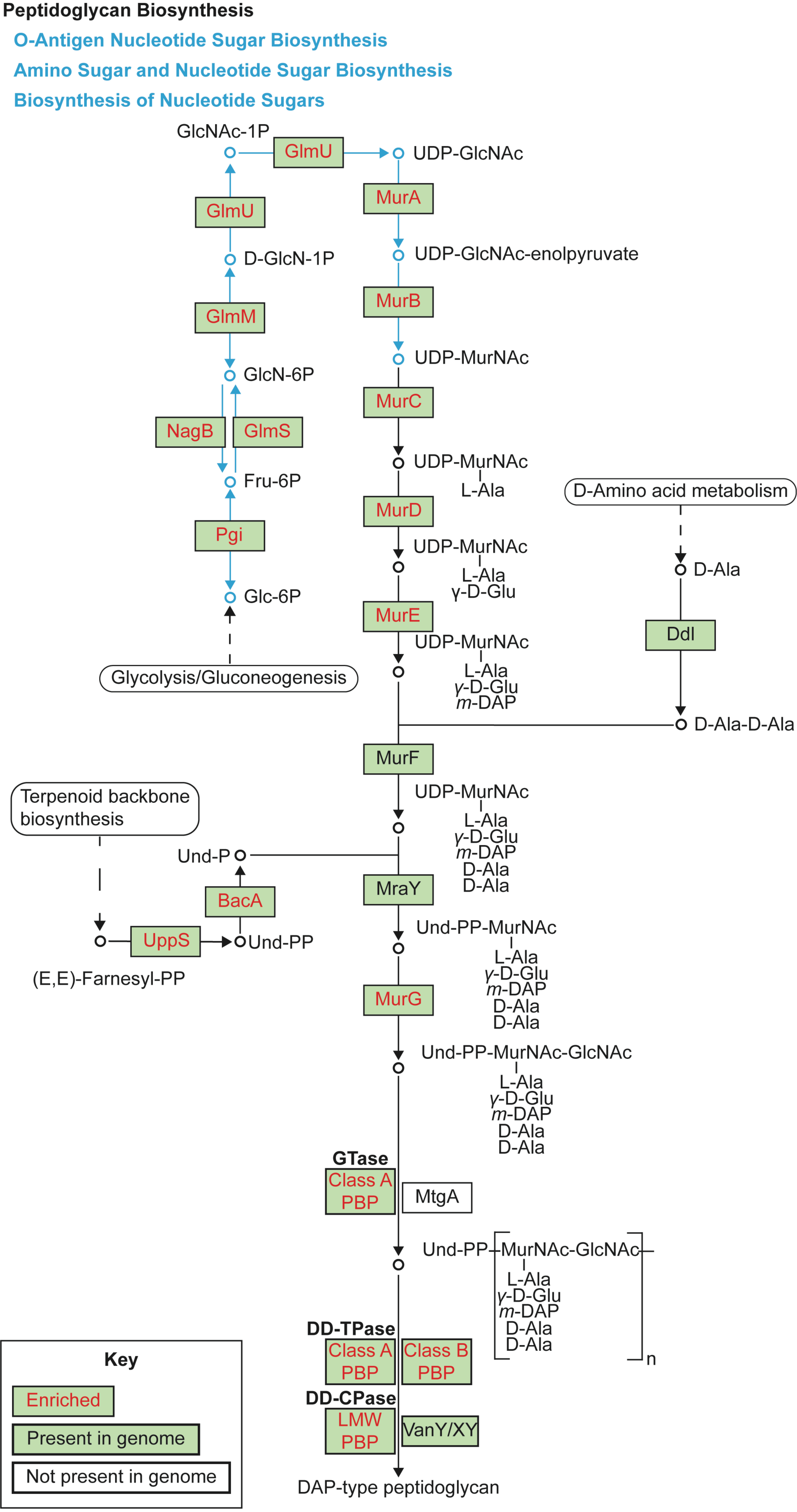


**Supplemental Figure S5.** Convergence of KEGG pathways most enriched in light.

The four KEGG pathways most enriched during light conditions are Peptidoglycan Biosynthesis, O-Antigen Nucleotide Sugar Biosynthesis, Amino Sugar and Nucleotide Sugar Biosynthesis, and Biosynthesis of Nucleotide Sugars. Enriched genes are presented in green boxes with red text, genes encoded in the Nostoc punctiforme but not enriched are presented in green boxes with black text, and genes not encoded in the N. punctiforme genome are in white unshaded boxes. Blue arrows indicate the biosynthetic pathways converging with peptidoglycan biosynthesis pathway (black arrows).
