## Supplemental Figure S6 for "Diel expression dynamics in filamentous cyanobacteria"

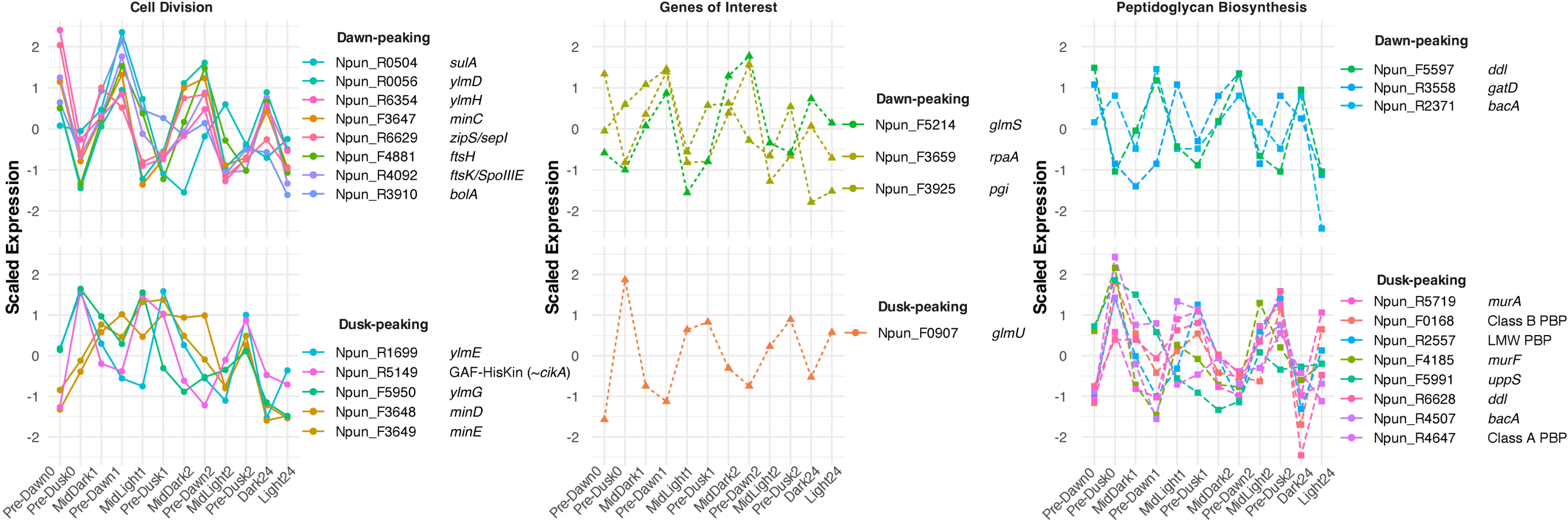


**Supplemental Figure S6.** Transcriptional convergence of cell division, sugar metabolism, and peptidoglycan biosynthesis.
Z-score normalized expression of genes involved in cell division, sugar metabolism, and peptidoglycan biosynthesis across different diel states. Expression values within each category are distinguished between dawn- and dusk-peaking patterns.
