## Supplemental Figure S7 for "Diel expression dynamics in filamentous cyanobacteria"

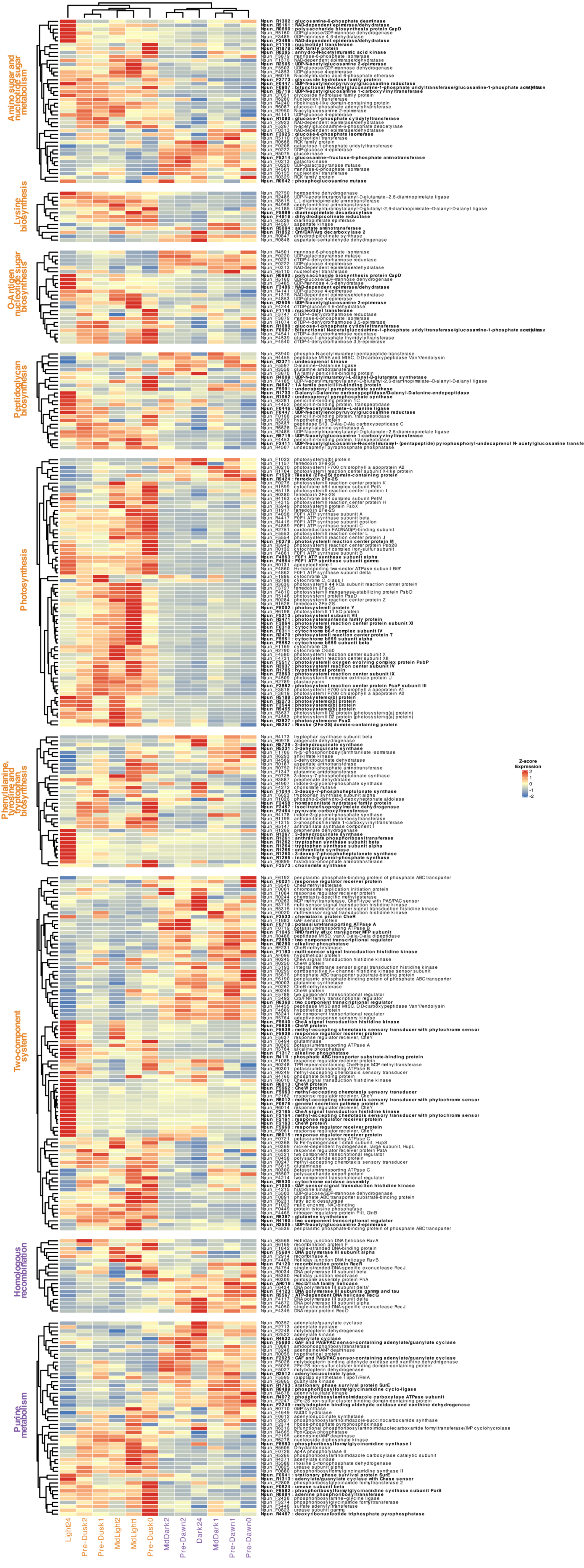


**Supplemental Figure S7.** Detailed Enriched KEGG Processes.

Hierarchical clustering (Euclidean distance, complete linkage) and z-score normalization (blue = downregulated, red = upregulated) of gene expression data highlight the light-associated (orange) and dark-associated (purple) enriched KEGG pathways. Enriched genes are bolded.
