## Supplemental Figure S8 for "Diel expression dynamics in filamentous cyanobacteria"

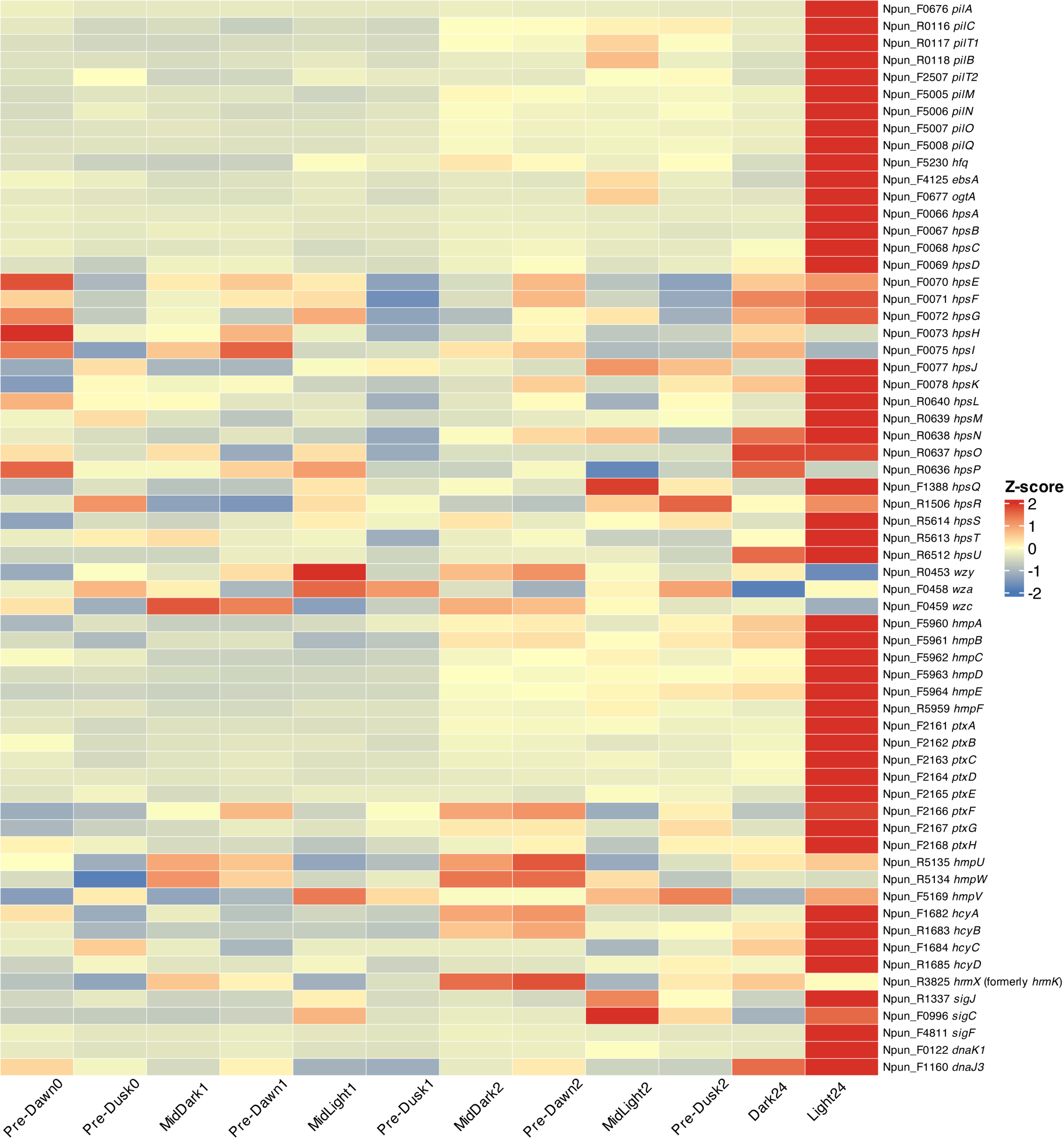


**Supplemental Figure S8.** Expression of hormogonium-associated genes.
Z-score normalized expression heatmap for all hormogonium-associated genes listed in Supplemental Table S6. Each row represents a gene, and each column represents a timepoint. The color gradient indicates the directional level of expression, with blue representing downregulation and red representing upregulation.
