## Supplemental Figure S9 for "Diel expression dynamics in filamentous cyanobacteria"

**Nucleotide Metabolism and Purine Metabolism**

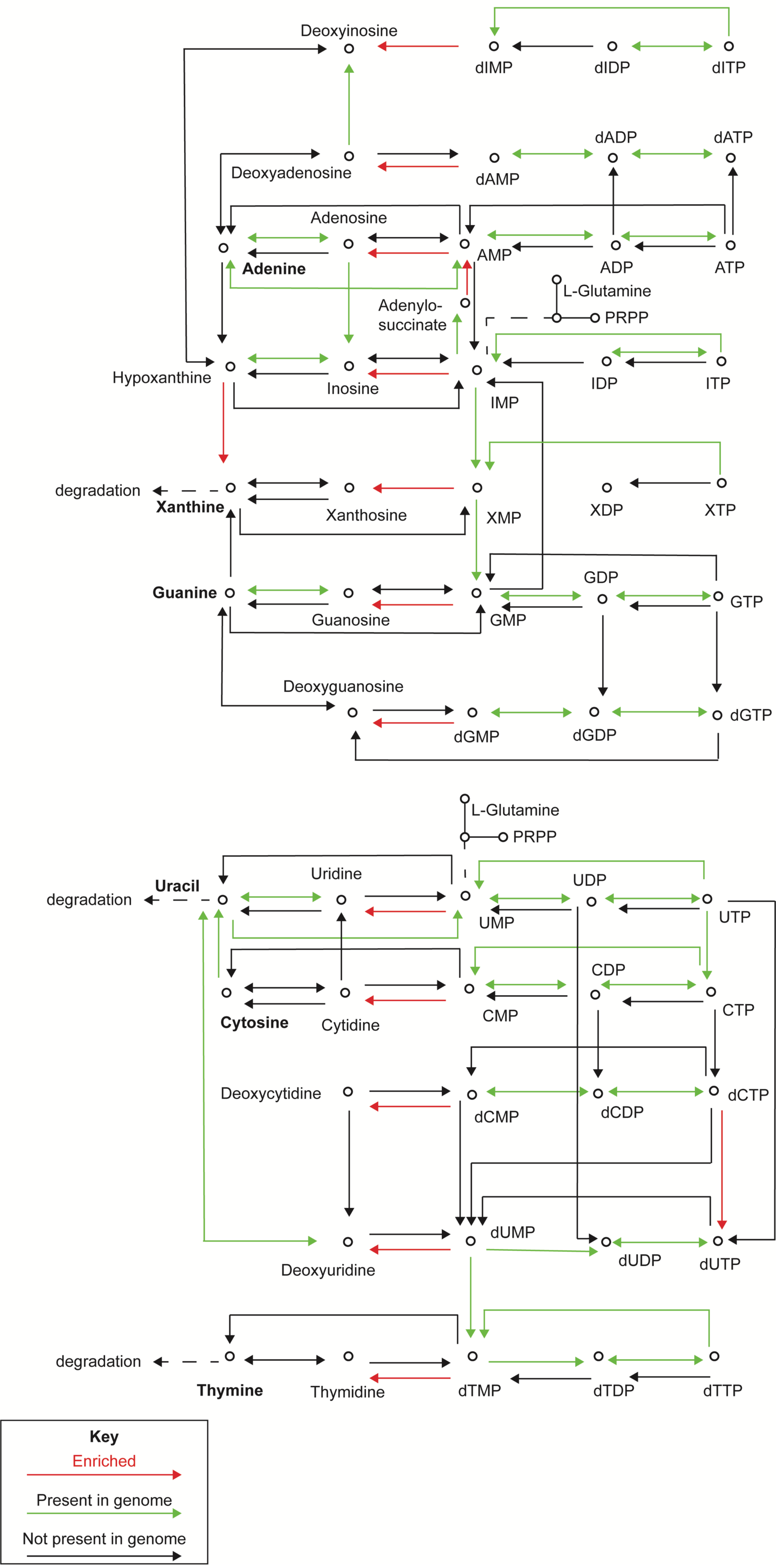


**Supplemental Figure S9.** Dark-associated nucleotide and purine metabolism pathways. Dark-associated pathways involved in nucleotide and purine metabolism, with red arrows indicating enriched genes, green arrows indicating pathways encoded in the genome but not enriched, and black arrows representing pathways not encoded in the genome.
