## Supplemental Figure S10 for "Diel expression dynamics in filamentous cyanobacteria"

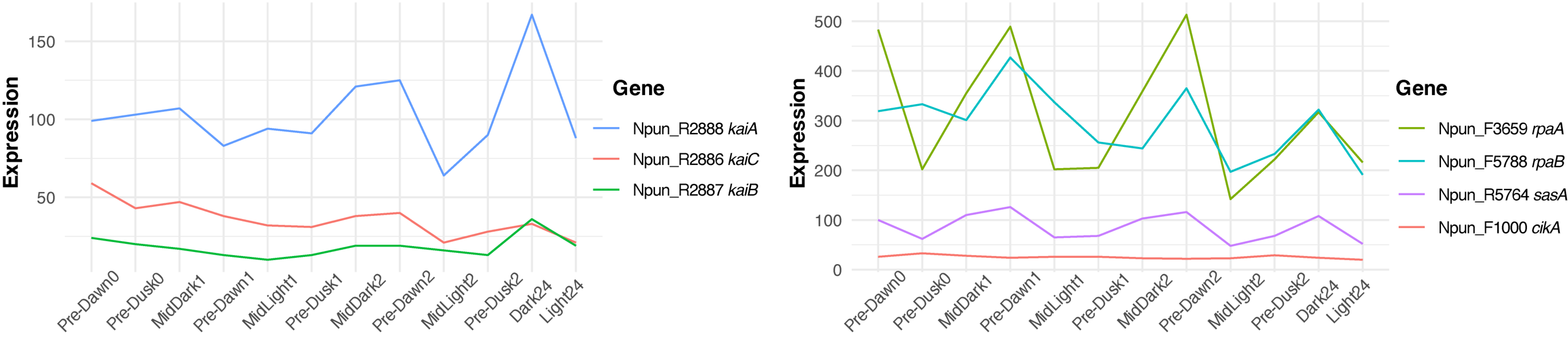


**Supplemental Figure S10.** Putative circadian gene expression.
(A) Expression levels of core Kai-protein clock genes (kaiA, kaiB, kaiC) and (B) non-core (rpaA, rpaB, sasA, cikA) across the time-course experiment.
