## Supplemental Figure S11 for "Diel expression dynamics in filamentous cyanobacteria"

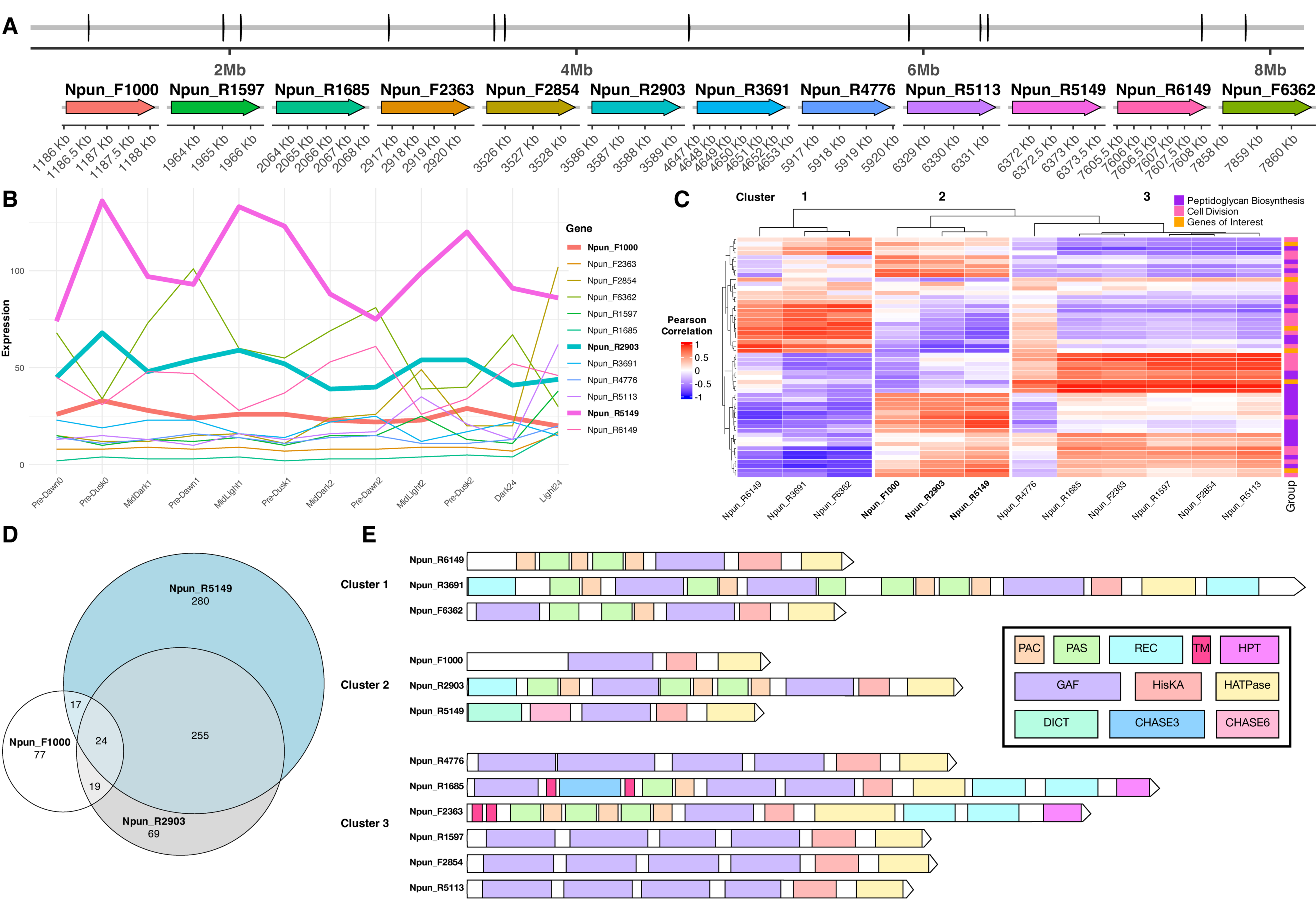


**Supplemental Figure S11.** N. punctiforme CikA-like paralogs containing GAF and histidine kinase domains.

(A) Genomic loci and nucleotide lengths of CikA-like paralogous genes in N. punctiforme. (B) Expression levels of CikA-like genes across the time-course experiment. (C) Hierarchical clustering (Euclidean distance, complete linkage) of Pearson correlations between CikA-like gene expression and selected genes within peptidoglycan biosynthesis (purple), cell division (pink), and key genes linking these functions (orange). The heatmap uses a blue-red gradient to indicate correlation values, with blue representing negative correlations and red representing positive correlations. (D) Euler diagram showing the distribution of global transcriptome genes with ≥0.75 correlation to the three CikA-like genes. (E) Domain architecture of the three clusters of 12 total CikA-like histidine kinases, highlighting the following domains: PAS (Per-Arnt-Sim), GAF (cGMP-specific phosphodiesterases, adenylyl cyclases, and FhlA), HisKA (Histidine Kinase A), HATPase (Histidine Kinase-Like ATPases), REC (Receiver domain), DICT (Diguanylate Cyclase domain), CHASE6 (Cyclase/Histidine kinase Associated Sensor domain 6), CHASE3 (Cyclase/Histidine kinase Associated Sensor domain 3), TM (Transmembrane domain), and HPT (Histidine Phosphotransfer domain).
