## Supplemental Figure S12 for "Diel expression dynamics in filamentous cyanobacteria"

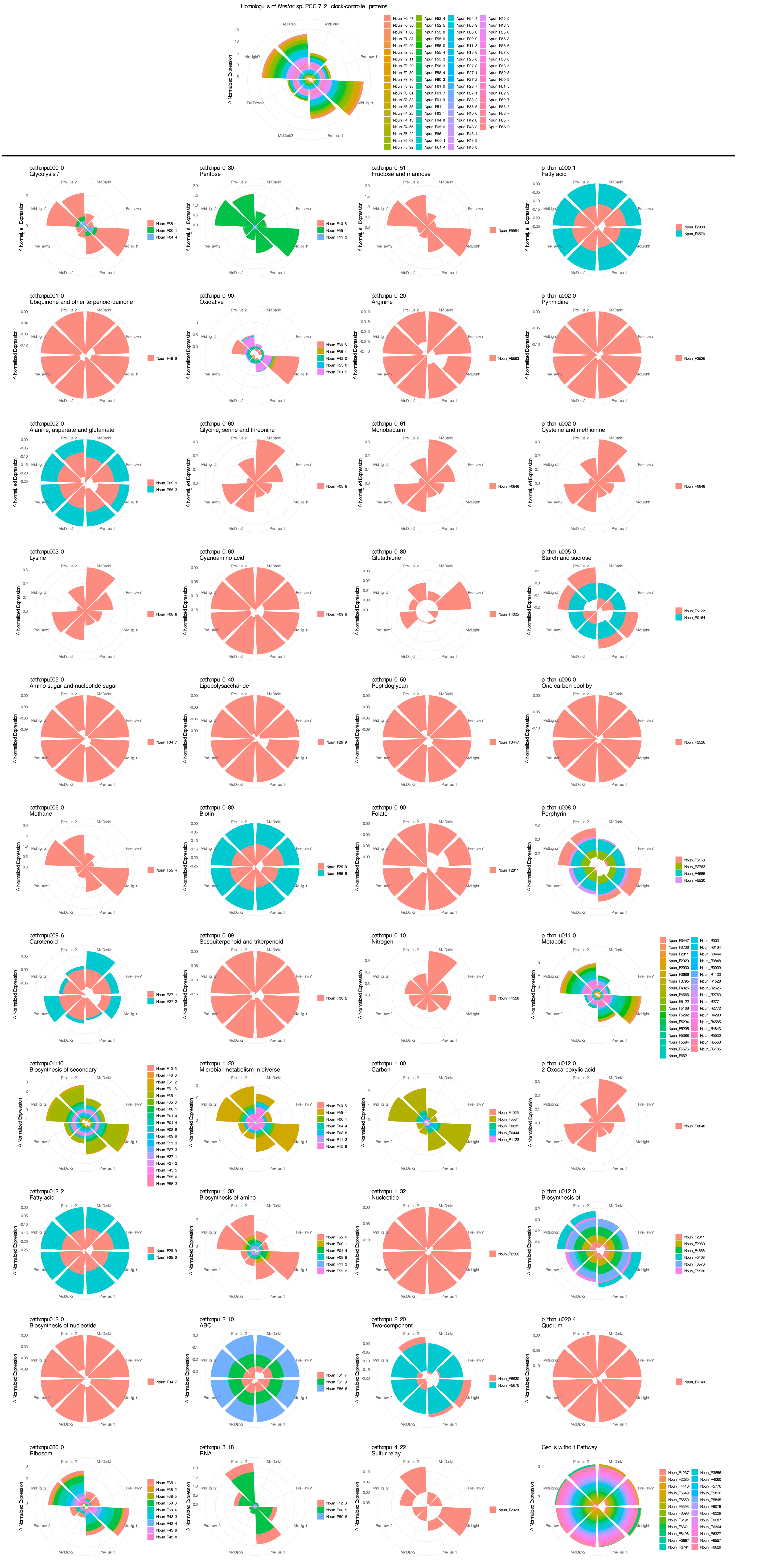

**Supplemental Figure S12.** Polar plots of clock-controlled protein homolog transcription.
Polar plots constructed by z-score normalizing each timepoint's transcriptome data for N. punctiforme genes homologous to Nostoc PCC 7120 experimentally-determined circadian clock-controlled proteins. (A) Polar plots excluding two highly expressed genes, cpcB (Npun_F5289) and cpcA (Npun_F5290), to maintain scale integrity. (B) Polar plots partitioned by KEGG pathways, showing the transcriptional patterns of clock-controlled protein homologues across diel states.
