## Supplemental Figure S13 for "Diel expression dynamics in filamentous cyanobacteria"

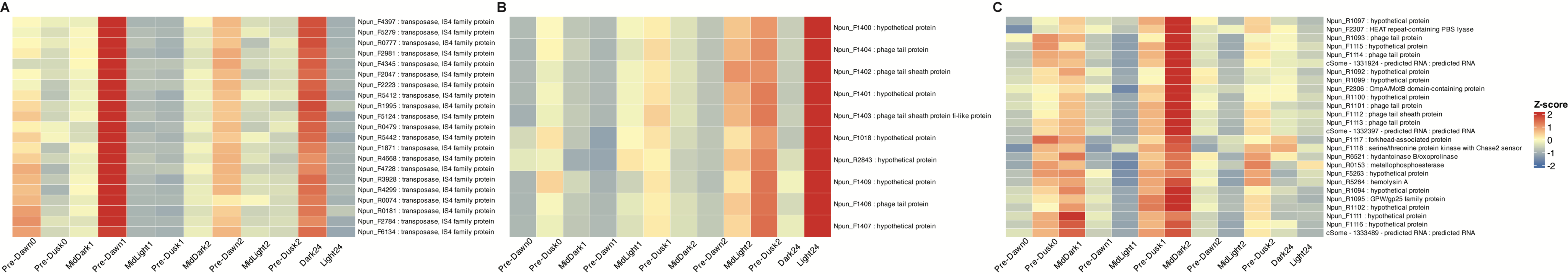


**Supplemental Figure S13.** Dynamically expressed mobile genetic elements.
Z-score normalized expression heatmaps of three dynamically expressed mobile elements across the time-course experiment, identified from the hierarchically clustered (Euclidean distance, complete linkage) transcriptome.
(A) Pre-Dawn1 + Dark24 cluster of IS4 family transposases. (B) Light24 cluster of phage proteins. (C) MidDark2 cluster of phage proteins. Heatmaps show expression patterns across the timepoints, with blue indicating downregulation and red indicating upregulation.
