## Supplemental Figure S14 for "Diel expression dynamics in filamentous cyanobacteria"

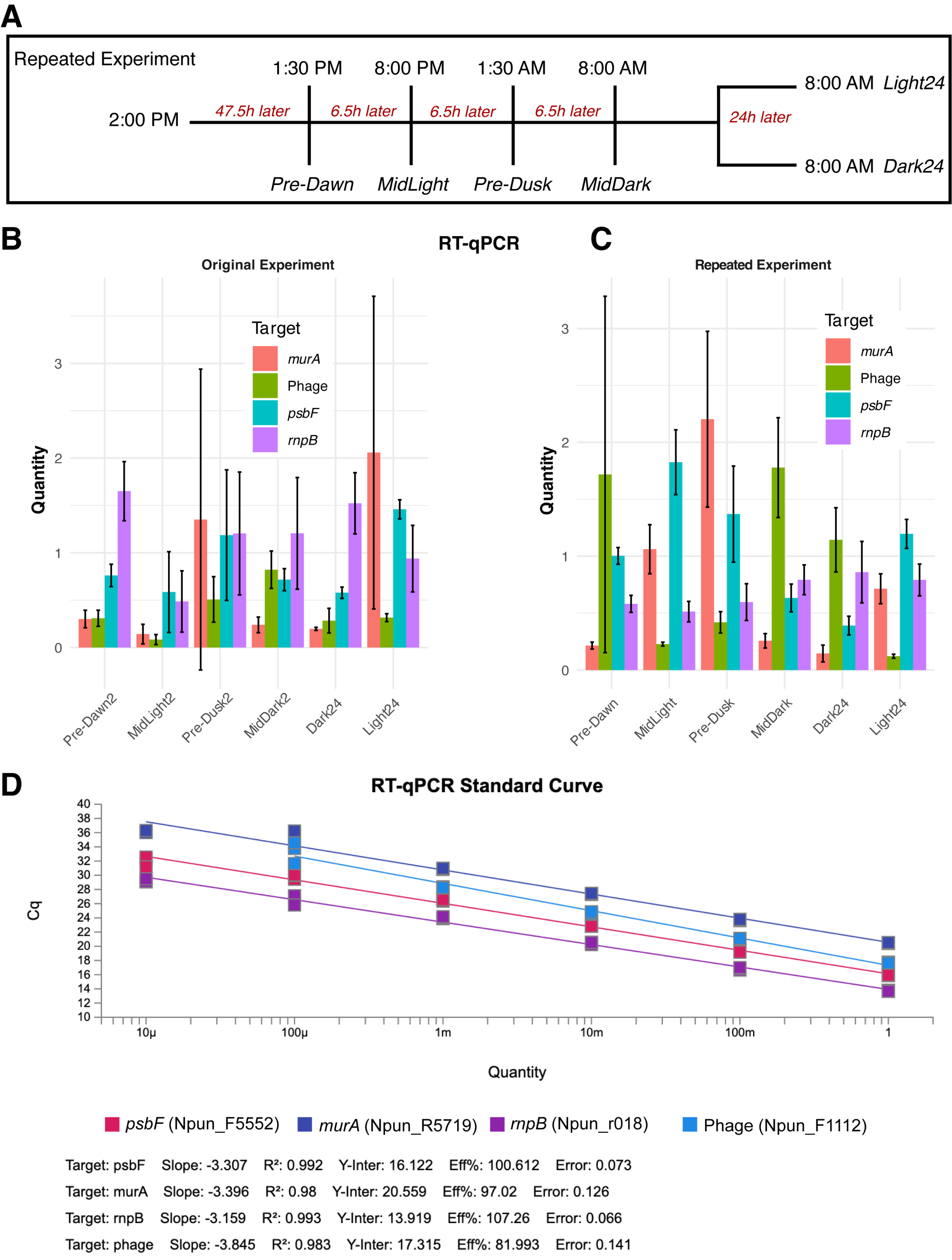


**Supplemental Figure S14.** Reverse transcription quantitative PCR analysis of gene expression.
(A) Repeated diel experiment collecting samples at Pre-Dawn, MidLight, Pre-Dusk, MidDark, and Light24 + Dark24 timepoints. (B) RT-qPCR absolute quantification of murA (DP-N-acetylglucosamine enolpyruvyl transferase), psbF (Cytochrome b559 beta subunit), and MidDark2 phage tail sheath protein, normalized to rnpB (RNase P RNA gene), for original experimental samples (Stages III and IV). (C) RT-qPCR absolute quantification for repeated experimental samples. (D) Standard curve for the RT-qPCR.
